# Nuclear Localization of Argonaute is affected by Cell Density and May Relieve Repression by microRNAs

**DOI:** 10.1101/2023.07.07.548119

**Authors:** Krystal C. Johnson, Audrius Kilikevicius, Cristina Hofman, Jiaxin Hu, Yang Liu, Selina Aguilar, Jon Graswich, Yi Han, Tao Wang, Jill M. Westcott, Rolf A. Brekken, Lan Peng, Georgios Karagkounis, David R. Corey

## Abstract

Argonaute protein is associated with post-transcriptional control of cytoplasmic gene expression through miRNA-induced silencing complexes (miRISC). Specific cellular and environmental conditions can trigger AGO protein to accumulate in the nucleus. Localization of AGO is central to understanding miRNA action, yet the consequences of AGO being in the nucleus are undefined. We show nuclear enrichment of AGO2 in HCT116 cells grown in two-dimensional culture to high density, HCT116 cells grown in three-dimensional tumor spheroid culture, and human colon tumors. The shift in localization of AGO2 from cytoplasm to nucleus de-represses cytoplasmic AGO2-eCLIP targets that were candidates for canonical regulation by miRISC. Constitutive nuclear localization of AGO2 using an engineered nuclear localization signal increases cell migration. Critical RNAi factors also affect the localization of AGO2. Knocking out an enzyme essential for miRNA biogenesis, DROSHA, depletes mature miRNAs and restricts AGO2 localization to the cytoplasm, while knocking out the miRISC scaffolding protein, TNRC6, results in nuclear localization of AGO2. These data suggest that AGO2 localization and miRNA activity can be regulated depending on environmental conditions, expression of mature miRNAs, and expression of miRISC cofactors. Localization and expression of core miRISC protein machinery should be considered when investigating the roles of miRNAs.

## INTRODUCTION

MicroRNAs (miRNAs) are short, ∼22 nucleotide regulatory non-coding RNAs that regulate gene expression through the RNA interference (RNAi) pathway (1,2). In mammalian cells, miRNAs are often associated with post-transcriptional repression of gene expression in the cytoplasm by the miRNA-induced silencing complex (miRISC). The miRISC complex is made up of an Argonaute (AGO) protein loaded with a miRNA guide. The miRNA carries the sequence-specificity to recognize a complementary messenger RNA (mRNA) target through Watson-Crick base pairing, and the AGO protein promotes recognition to silence the expression of this target directly by enzymatic cleavage or indirectly by recruiting other miRISC effector proteins to inhibit translation and/or accelerate the degradation of the mRNA (3-6). There are four AGO variants in human cells, AGO1-4. AGO2 (3,4,5), and to a lesser extent AGO3 (7), can promote cleavage of target transcripts.

miRNAs also have the potential to function in the nucleus of human cells. Both miRNAs (8), AGO proteins (8-13), and other miRISC cofactors can be detected in cell nuclei and chromatin (11,14,15). The relative localization of AGO between cytoplasm and nuclei varies in different cell types and tissues (16). Synthetic small RNAs and miRNAs have been shown to control gene transcription and splicing (17-32), demonstrating the potential for small RNAs to control nuclear gene expression.

While exogenous duplex RNAs can control gene expression in mammalian cell nuclei, the importance of the RISC machinery shuttling to the nucleus for regulation of endogenous gene expression has not been well-established. Our laboratory has previously used CRISPR/Cas9-mediated knockout cells deficient for the primary AGO variants AGO1, AGO2, and AGO3 to examine the scope of RISC-mediated regulation in the cytoplasm and nuclei of HCT116 colon cancer-derived cells (33,34). When these cells were grown under normal cell culture conditions, we found that knocking out the expression of AGO protein produced modest effects on gene expression relative to wild-type cells. This observation is consistent with previous observations that, despite large numbers of target genes predicted to be regulated by miRNAs, most single miRNA knockout animals show no apparent phenotype until placed under stress (35-40). We hypothesized that specific environmental conditions might enhance the impact of RNAi, affect the subcellular localization of the miRISC machinery, and provide greater insights into the endogenous regulation of miRNA activity.

Other laboratories have previously suggested that specific environmental conditions might change the localization of AGO and alter the relative impact of RISC-mediated regulation in the cytoplasm and nucleus. In 2011, Sczakiel and coworkers observed that varied cellular stresses, including heat shock, sodium arsenite, cycloheximide, and lipid-mediated transfection of phosphorothioate oligonucleotides could change the sub-cellular localization of AGO2 protein (41). This study suggested that these stressors promote inclusion of AGO2 in stress granules leading to a reduction in cytoplasmic RNAi activity. Since then, other studies have linked cell stress to changes in the relative localization of AGO protein between nuclei and cytoplasm (42). Nuclear translocation of AGO2 can be induced by senescence (43,44) and has been suggested to be a trigger for miRNA-mediated transcriptional gene silencing (43). Nuclear AGO2 has recently been reported to silence mobile transposons in quiescent cells (45) and enhance expression of genes in spermatogenic cells through association with meiotic chromatin (14).

Here we examine the effect of cell density, cell contact, and tumor formation on the subcellular localization of AGO protein and miRNA regulation. We observe that in two-dimensional cell culture grown to high density, three-dimensional tumor spheroid cell culture, or human tissue samples, nuclear localization of AGO2 increases. In cell culture, this relocalization de-represses the expression of AGO2-bound genes that are candidates for canonical silencing by miRNAs in the cytoplasm. These data suggest that miRNA regulation and the subcellular localization of AGO2 are dynamic systems that are sensitive to environmental change, the availability of miRNAs and miRISC cofactors, and tumor formation.

## MATERIALS AND METHODS

### Tissue Culture

Wild-type HCT116, DROSHA-/-, TNRC6AB-/-, AGO1/2/3-/- and NLS-AGO2 KI cell lines were cultured in McCoy’s 5A medium supplemented with 10% FBS. A549 cells were grown in F12K medium supplemented with 10% FBS. HeLa cells were cultured in Dulbecco’s Modified Eagle *Medium* (*DMEM*) with 10% FBS. HepG2 were grown in Eagle’s Minimal Essential Medium (EMEM) with 10% FBS. Healthy control fibroblast cell line (GM02153) was purchased from Coriell institute, which was cultured in Minimal Essential Medium Eagle (MEM) with 10% FBS. All cells were incubated at 37°C in 5% CO_2_ and passed when 70-80% confluent.

### Preparation of cytoplasmic and nuclear extract from cell culture

Cells were seeded at 100,000 cells per well into 6-well plates on Day 0, and 8 mL of fresh media was supplied every day starting on Day 2 through Day 7. Cells were detached with trypsin, harvested, and counted with trypan blue staining (TC20 Automated Cell Counter; Bio-Rad). Subcellular fractionation to isolate cytoplasm and nuclear lysate extract was similar as previously described (11,12,33,34).

### Preparation of cytoplasm, nuclear, and chromatin extract from patient tissue

Patient tissue was kept frozen with liquid nitrogen and cut into small pieces with a blade. The small tissue pieces were homogenized by crushing in a mortar and pestle with liquid nitrogen replenished to keep tissue frozen. Homogenized tissue was fractionated per the manufacturer’s protocol using the Subcellular Protein Fractionation Kit for Tissues (ThermoFisher, Cat No. 87790).

### Dual Viability Staining Assay for Apoptosis Detection with Flow Cytometry

Invitrogen™ eBioscience™ Annexin V Apoptosis Detection Kit FITC (Cat. No. 88-8005-72) was used in combination with Propidium Iodide Solution (Sigma-Aldrich, Cat. No. P-4864-10mL) as described in manufacturer’s protocol. Cells were analyzed by flow cytometry, and summary plots were generated with FlowJo software. Three biological replicates were performed for each cell density collection time point.

### RNA half-life assay

Cells were seeded at 100K cells per well in 6-well plates on Day 0. At 50% (Day 3) and 400% (Day 7) confluence, the growth media was replaced with growth media supplemented with 10 µg/ml of Actinomycin D (Sigma Aldrich: A4262-5MG). Cells were collected at time points: 0, 1, 2, 4, 8, 12, 18, 24, 36, 48 hours and then fractionated with cytoplasmic and nucleus fractionation protocol. The total RNA from cytoplasmic fraction was isolated using TRizol LS reagent, nuclear pellet – TRizol reagent. Data points were kept uniform by using exactly 2 µg of RNA pretreated with DNase as input in reverse transcription reaction. RNA stability was measured between 50% (Day 3) and 400% (Day 7) of seven different potential reference genes to analyze stability assay data. The input of cDNA synthesis was normalized across samples, and the data was analyzed by differences in stability based on changes in Ct value. All reference genes and experimental genes were tested from the same cDNA. Pre-mRNA levels were checked in the nuclear fraction of cells harvested at 50% and 400% to validate that transcription was inhibited at the same time by Actinomycin D treatment in both density conditions.

### Co-Immunoprecipitation

Cells were lysed with whole cell lysis buffer (1xPBS, Igepal 0.5%, Na-deoxycholate 0.5%, SDS 0.1%, 1x Protease inhibitors), supplemented with 10% glycerol and snap frozen before immunoprecipitation. 335 µg of total protein was incubated with 80 µL Dynabeads Protein G (Invitrogen: 10004D), and 3 µg of anti-AGO2 polyclonal antibody. This antibody was generated by GenScript, and is derived from the same peptide sequence used to elicit “3148” from the Nelson’s lab (46). We also use antibodies against TNRC6A (Bethyl: A302-329A) or species matching IgG antibody (Millipore:) in 1 mL of lysis buffer and incubate with rotation over night at 4 °C. Wash six times with 1 mL of immunoprecipitation wash buffer (Tris-HCl, PH-7.4 20 mM, NaCl 400 mM, MgCl_2_ 3 mM, NP-40 0.05%, SDS 0.01%) with rotation for 5 min at 4 °C. The protein was eluted with 20 µl of 2x Tris-SDS loading buffer incubating for 20 min at room temperature. The elute was transferred to fresh tube heated for 10 min at 95 °C and ran on 4-20% SD-PAGE gel. The western-blot was performed under the same condition in this paper using following antibodies: anti-AGO2 (WACO: 015-22031), anti-TNRC6A (Bethyl: A302-329A).

### 3D Tumor Spheroid Culture

Single cells were resuspended in a 40 µL mixture of 70% activated rat tail Collagen 1 (Corning, 354236) and 30% Cultrex UltiMatrix Basement Membrane Extract (R&D systems, BME001) at a final concentration of 2.1 mg/mL Collagen 1 and 3.5mg/ml UltiMatrix BME. The embedded cells were plated in 8-well chamber slides (BD Biosciences) and allowed to gel at 37°C. Each well of the chamber slide contained 2,000 cells. After 1 hour, cultures were supplemented with DMEM media (Corning, 10013CV) containing 10% FBS and 1% Penicillin-Streptomycin (Gibco, 15140122). Cultures were incubated for 7 days at 37°C and 5% CO_2_, with fresh media added on day 4. After 7-8 days of growth, the ECM was dissolved with dispase (Stemcell Technologies, 07913), cells were washed 2X with PBS, and then lysed. Cultures were imaged at day 7 (prior to dispase treatment) at 10X magnification. Scale bars = 50 µm.

### Western-blot analysis

Total protein lysate was prepared re-suspending cells in lysis buffer (50 mM Tris–HCl, pH 7.0, 120 mM NaCl, 0.5% NP-40, 1 mM EDTA, 1 mM DTT, 1× protease inhibitor (Roche, complete). Proteins were separated on 4–20% gradient Mini-PROTEAN® TGXTM precast gels (Bio-Rad). After gel electrophoresis, proteins were wet transferred to nitrocellulose membrane (0.45 μM, GE Healthcare Life Sciences) at 100 V for 75 min. Membranes were blocked for 1 h at room temperature with 5% milk in 1× PBS containing 0.05% TWEEN-20. Blocked membranes were incubated with the primary antibodies in blocking buffer at 4°C on rocking platform overnight: using anti-AGO1, 1:2000 (5053, Cell Signaling), anti-AGO2, 1:1500 (015-22031, Fujifilm WAKO), anti-Calnexin, 1:1000 (2433, Cell Signaling), anti-Lamin A/C, 1:1500 (ab8984, Abcam), anti-β-actin, 1:5000 (A5441, Sigma-Aldrich), anti-TNRC6A (GW182), 1:5000 (A302-329A, Bethyl), anti-β-tubulin, 1:5000 (32-2600, Invitrogen) antibodies. After primary antibody incubation, membranes were washed 3 × 10 min at room temperature with 1× PBS+0.05% Tween-20 (PBST 0.05%) and then incubated for 1 h at room temperature with respective secondary antibodies in blocking buffer. Membranes were washed again 4 × 10 min in PBST 0.05%. Washed membranes were soaked with HRP substrate according to manufacturer’s recommendations (SuperSignal™ West Pico Chemiluminescent substrate, Thermo Scientific) and exposed to films. The films were scanned and bands were quantified using ImageJ software.

### CRISPR/Cas9 Generation of NLS-AGO2 HCT116

The SV40 large T-antigen NLS sequence that was previously reported to manipulate AGO2 localization (13,47) (**Supplemental Figure 13**) was inserted into the N-terminus of human AGO2 using CRISPR/cas9 in WT HCT116 cells carried out by GenScript. The SV40 NLS amino acid sequence is PKKKRKVAG, and the DNA sequence is CCAAAAAAGAAGAGAAAGGTAGCTGGT. NLS fragment had 2 amino acids at both side as spacer sequences. NLS was inserted into both alleles at Exon 1 to force AGO2 localization to the nucleus. To maintain endogenous transcription level, the endogenous promoter was used.

### NLS-AGO2 HCT116 vs WT HCT116 Growth Curve

Cells were grown in six well plates. Cells were seeded on Day 0 at a concentration of 1×10^5^ cells per well. Cells were harvested and counted daily, from Day 1 to Day 7. To count the cells, cells were mixed with equal volumes of trypan blue and were counted using a cell counter (TC20 Automated Cell Counter; Bio-Rad). Cell media was replenished daily, starting on Day 2, to avoid pH stress and nutrition deprivation.

### Argonaute 2 absolute protein quantification

Quantification was completed as described (48) with modifications. Briefly, recombinant AGO2 protein (Active Motif) was purchased and serially diluted in Protein LoBind tubes (Eppendorf) coated with Bovine Serum Albumin standard protein (ThermoFisher). Cells were collected at low (Day 3) and high confluence (Day 7) time points and subcellular fractions were isolated as in Cytoplasm/Nuclear Fractionation. Fractions (whole cell, cytoplasm, and nucleus) were lysed to contain 10,000 cells/µL based on initial cell count from collection. Recombinant protein standard curves were resolved by western blot and analyzed with ImageJ to construct a standard curve. The slope of the curve was used to determine the AGO2 proteins per cell in whole cell, cytoplasmic and nuclear lysates. Statistical analysis was performed in Prism using an unpaired t-test with Welch’s correction.

### AGO2-GFP plasmid expression in AGO1/2/3-/- cells

AGO1/2/3 knockout (AGO1/2/3-/-) cells previously described in Chu et al., 2020. The WT-AGO2-GFP and NES-AGO2-GFP plasmids were generated by GenScript. A classic leucine-rich nuclear export signal (NES), LAQQFEQLSV, was used from *Arabidopsis thaliana* AGO1 (49) and in the NES database (http://prodata.swmed.edu/LRNes/IndexFiles/details.php?name=313). The human AGO2 (WT-AGO2, NCBI Reference Sequence: NM_012154.5) and the N-terminal NES tagged AGO2 (NES-AGO2) were cloned into mammalian expression vector pcDNA3.1+N-eGFP. AGO1/2/3-/- cells were transfected with WT-AGO2-GFP or NES-AGO2-GFP plasmids with OptiMEM and FuGENE HD Transfection Reagent (Promega, Cat. No. E2311) with media change to full media at 24 hours post-transfection. Cells were detached with trypsin and collected 48 hours post-transfection. Cells were washed with PBS and sorted for GFP-positive cells using FACS. Sorted cells were diluted in PBS, then used for (1) whole cell RNA extraction, (2) subcellular protein fractionation, and (3) adhered to microscope slides using a Cytospin cytocentrifuge. The cells used for microscopy were fixed with 4% paraformaldehyde for 10 min, washed with PBS and permeabilized for 10 min with 0.2% Triton X-100 in PBS. Fixed and permeabilized cells were washed with PBS. Then, Vectashield Hard Set mounting medium with DAPI was added and covered with a coverslip. After the mounting medium was set, cells were imaged with a 60X objective lens and 405 nm and 488 nm lasers on a Nikon CSU-W1 with SoRa spinning disk confocal microscope. Images were then processed in ImageJ.

### Transfection of duplex RNA

All duplex RNA transfections used Lipofectamine RNAi MAX (Invitrogen). 24 hours before transfection, cells were seeded into six-well plates at 150K cells per well for wild type HCT116 cells and 300K for DROSHA-/- cells. At the next day, 50 nM of siRNAs were transfected into cells as previously described. (50) The transfected siRNAs included siCM as a negative control, an siRNA targeting MALAT1 as described in Liu et al. 2019, and duplex miRNA mimics of let-7a and miR-27a. 24 hours later, cells were replaced with full culture media. Cells were harvested for western blot 48 hours after transfection.

### RT-qPCR for candidate genes

Total RNA was extracted from cells and treated with DNase I (Worthington Biochemical) at 25°C for 20 min, 75°C for 10 min. Reverse transcription was performed using high-capacity reverse transcription kit (Applied Biosystems) per the manufacturer’s protocol. 2.0 μg of total RNA was used per 20 μL of reaction mixture. PCR was performed on a 7500 real-time PCR system (Applied Biosystems) using iTaq SYBR Green Supermix (BioRad). PCR reactions were done in triplicates at 55°C 2 min, 95°C 3 min and 95°C 30 s, 60°C 30 s for 40 cycles in an optical 96-well plate. For each gene, two different sets of primer were used to check mRNA level (Supplementary Table S1). Data were normalized relative to internal control genes HPRT1, snRNP200, and ZMYM4 levels.

### RT-qPCR for miRNAs

Total RNA was extracted from cells with TRIzol as described above. RNA from cytoplasm lysate was extracted with TRIzol LS (Invitrogen, Cat. No. 10296028) and from nuclear pellet with TRIzol. The miRNAs were reverse transcribed using universal miRCURY LNA kit (Qiagen: 339340). The cDNA was further used for quantification of miRNAs following the protocol of miRCURY LNA SYBR Green kit (Qiagen: 339346). PCR was performed on a 7500 real-time PCR system (Applied Biosystems) using TaqMan Universal PCR Master Mix No AmpErase UNG (Applied Biosystems) with the primers and probes included in the TaqMan microRNA Assay kits (ThermoFisher, hsa-miR-210-3p: Assay ID: 000512; hsa-miR-181a-5p: Assay ID: 000480; hsa-miR-21-5p: Assay ID: 000397; hsa-miR-27a-3p: Assay ID: 000408). PCR reactions were done in triplicates at 55°C 2 min, 95°C 3 min and 95°C 30 s, 60°C 30 s for 40 cycles in an optical 96-well plate.

### Preparation of whole transcriptome RNAseq libraries

Total RNA was extracted from whole cells grown in 2D culture for 3 days (50% confluency), 5 days (250% confluency), and 7 days (300-400% confluency) with TRIzol as described above. The RNA quality was measured using the Agilent 2100 Bioanalyzer system with the RNA Nano chip kit (catalog# 5067-1511), and the RNA concentration was measured using the Nanodrop 2000 spectrophotometer by Thermo Scientific. Roche Kapa RNA HyperPrep Kits with RiboErase (HMR) (Catalog #KK8561) were used to generate the RNA libraries. The procedure was followed with the manufacturer’s instructions. The RNA input amount for each library preparation was 1.0 µg. The workflow began with rRNA depletion, followed by RNAse H and DNase treatment. Subsequently, fragmentation was carried out at 94°C for 5 minutes. The cleaved RNA fragments were then reverse transcribed into first strand cDNA synthesis using reverse transcriptase and random primers. Second strand cDNA synthesis and A-tailing were performed, with incorporation of dUTP into the second cDNA strand. UMI adapters synthesized by IDT (Integrated DNA Technology) were used during the ligation process. The cDNA fragments underwent purification and enrichment via 9 to 10 cycles of PCR amplification to create the final cDNA library. Library quality was verified using the Agilent 2100 Bioanalyzer, and the concentration was measured using the picogreen method. The nine RNA-seq libraries were sequenced on Illumina NovaSeq 6000 sequencer platform using the PE-150 bp (paired-end) protocol. Image analysis and base calling were performed using an Illumina pipeline with default settings.

### Whole transcriptome RNAseq analysis of mRNA expression

Paired-end demultiplexed fastq files were generated using bcl2fastq2 (Illumina, v2.17), from NovaSeq6000 SP4 reagent’s bcl files. Initial quality control was performed using FastQC v0.11.8 and multiqc v1.7. Fastq files were imported batch wise, trimmed for adapter sequences followed by quality trimming using CLC Genomics Workbench (CLC Bio, v23.0.3). The imported high-quality reads were mapped against gene regions and transcripts annotated by ENSEMBL v99 hg38 using the RNA-Seq Analysis tool v2.7 (CLC Genomics Workbench), only matches to the reverse strand of the genes were accepted (using the strand-specific reverse option). Differential gene expression analysis between the sample groups was done using the Differential Expression for RNA-seq tool v2.8, where the normalization method was set as TMM. Differential expression between the groups was tested using a control group, outliers were downweighed, and filter on average expression for FDR correction was enabled.

### RNA extraction and small-RNA-seq analysis of miRNA expression

Whole cell RNA was extracted from cells seeded at 100K/well in 6-well plates on Day 0 and grown for 3 days (50% confluency), 5 days (250% confluency), and 7 days (300- 400% confluency) with Trizol. This whole cell RNA was submitted to the UTSW Genomics Sequencing Core for small-RNA-sequencing (also referred to as microRNA-sequencing). The miRNA libraries were prepared using Illumina TruSeq Small RNA library preparation kits (Catalog # RS-200-0012, RS-200-0024). Protocol from TruSeq® Small RNA Library Prep Reference Guide (Document # 15004197 v02, July 2016) was followed. The total RNA quality was checked with Bioanalyzer (Agilent, RNA 6000 Nano kit, 5067-1511). Small RNA library preparation started with 1ug high quality total RNA. First, the small RNAs were ligated with 3’ Adapter and then ligated with 5’ Adapter. Reverse transcription followed by amplification creates cDNA constructs based on the small RNA ligated with 3’ and 5’ adapters. Then libraries were cleaned with Agencourt AMPure beads (Beckman Coulter, Catalog # A63882). Then the concentration of the remaining clean RNA was measured by Picogreen (Fisher scientific, Quant-iT PicoGreen dsDNA Assay Kit, P7589). Equal amounts were pooled from each library. Then the library pool was further purified using Pippin Prep system (Sage Science INC) with 3% agarose gel cassettes (Sage Science INC, CDP3010). The final library pool quality was checked with Bioanalyzer (Agilent, High Sensitivity DNA Kit, 5067-4626) and qPCR (Kapa library quantification kit, 7960336001). Pooled libraries were sequenced on NextSeq v2.5 High Output flow cells as single end 50 cycle runs, the yield per library was between 6.9- 16.3 million pass filter reads, with a mean quality score of 34.6 and a % >= Q30 of 95.8% bases. Cutadapt trimmed read lengths were between 25 to 40 bases, mapping reference was mature and hairpin miRNA sequences from GRCh37. The same library preparation methods were used for all samples to obtain both hairpin and mature reads. The data were generated by trimmed mean of M-values method (TMM) normalized counts (51,52) that aligned with miRNA precursors or mature miRNAs. The data was part of the package generated by edgeR.

### Exon-Intron Split Analysis

Exon-Intron Split Analysis (EISA) was used to measure changes in mature and precursor mRNA reads and differentiate transcriptional from post-transcriptional regulation between different experiment groups (53). eisaR v1.2.0 package2 was employed to do the EISA between Day 3 (50%) versus Day 5 (250%) and Day 3 (50%) versus Day 7 (400%). Genes with FDR< 0.001 were defined as significantly post-transcriptionally regulated. Default parameters were employed.

### Pathway Analysis

The functional enrichment analysis was performed using g:Profiler (version e109_eg56_p17_1d3191d) with g:SCS multiple testing correction method applying significance threshold of 0.05 (54).

### Gene Network Analysis

Gene coexpression networks analysis was conducted with BioNERO V1.2.03 (55). Missing values in gene expression data were replaced with 0. Non-expressed genes whose median value was below 5 were removed. Top 8,000 most variable genes were kept. Confounding artifacts were adjusted with PC_correction function in BioNERO. Then the gene coexpression network was inferred according to the official pipeline of BioNERO.

### Scratch/Wound Healing/Migration Assay

This assay was performed as described (56). Briefly, HCT116 WT and NLS-AGO2 cells were seeded at 300,000 cells per well in 6-well plates to achieve 80-100% confluence 24 hours later. A 1 mL pipette was used to create a scratch in each well. Wells were imaged in the same location and orientation at 4X and 10X magnification every 12 hours for a duration of 96 hours. Three biological replicates at all time points analyzed for each cell line, and data was analyzed in ImageJ (NIH) as described (57) to calculate migration rate.

### Immunfluorescence microscopy in HCT116 WT and DROSHA-/-

HCT116 WT and DROSHA-/- cells were harvested from 6-well plates at desired confluency and prepared as a single cell suspension in cold PBS. The cells were spun down onto the slides using cytospin at low speed. Cells were fixed with 4% paraformaldehyde for 15 min at RT and then washed three times with 1X PBS. Cells were permeabilized with 0.3% Triton X-100 in 1X PBS and incubated at RT for 30 min. Then the cells were washed three times with 1 X PBS. Then cells were blocked with 5% normal goat serum in 1X PBS for 60 min at RT. After blocking step, the cells were washed three times with 1X PBS. Primary antibodies of anti-human AGO2 (Fujifilm Wako, clone: 4G8) were diluted with 1% BSA in 1 X PBS, incubated at 4°C overnight, and then cells were washed three times with 1X PBS. Secondary antibodies of goat-anti-mouse IgG (H+L) Alexa Fluor 488 (ThermoFisher, A 11001) were diluted 1:1000 with 1% BSA in PBS and incubated at 4°C for 60 min. The cells were washed with 1X PBS three times and mounted with VECTASHIELD® HardSet™ Antifade Mounting Medium with DAPI (Vector Laboratories, H-1500-10). The slide images were taken with Spinning Disk Confocal Nikon CSU-W1-SoRA microscope with 100x /1.49 oil SR HP Apo-Tirf objective.

## RESULTS

### AGO2 shifts localization to the nucleus in cells grown to high density

Cancer cells within a solid tumor must survive harsh environmental conditions such as hypoxic cores, overcrowding, and expansion to new environments for metastases (38). Growing cells to high density to form several overcrowded layers provides a simple cancer model system to examine the effect of changing environmental conditions on RISC factors. We chose to focus our study on HCT116 cells, which are a patient-derived colorectal cancer line that will form a tumor capable of metastasis when xenografted in mice. HCT116 cells are also diploid, making it relatively straightforward to obtain knockout and knock in cell lines (33). Counting cells grown within a defined surface area provides a quantitative measure of cell density, and this was used to establish an estimated baseline of how many cells were required to form a complete monolayer within the cell culture well, which we refer to as 100% confluent.

We grew HCT116 cells for seven days to reach a calculated confluence of 370% (approximately four stacked layers of cells) in two-dimensional cell culture (**Figure 1A**) and then fractionated the cells to determine the subcellular localization of AGO2, the most abundant AGO paralog in wild-type HCT116 cells (50). Confocal microscopy revealed a shift in the localization of AGO2 to the nucleus when cells were grown to 300-400% confluence (**Figure 1BC**). Western analysis confirmed the increased nuclear localization of AGO2 in cells grown to high density (**Figure 1D**).

**Figure 1.**
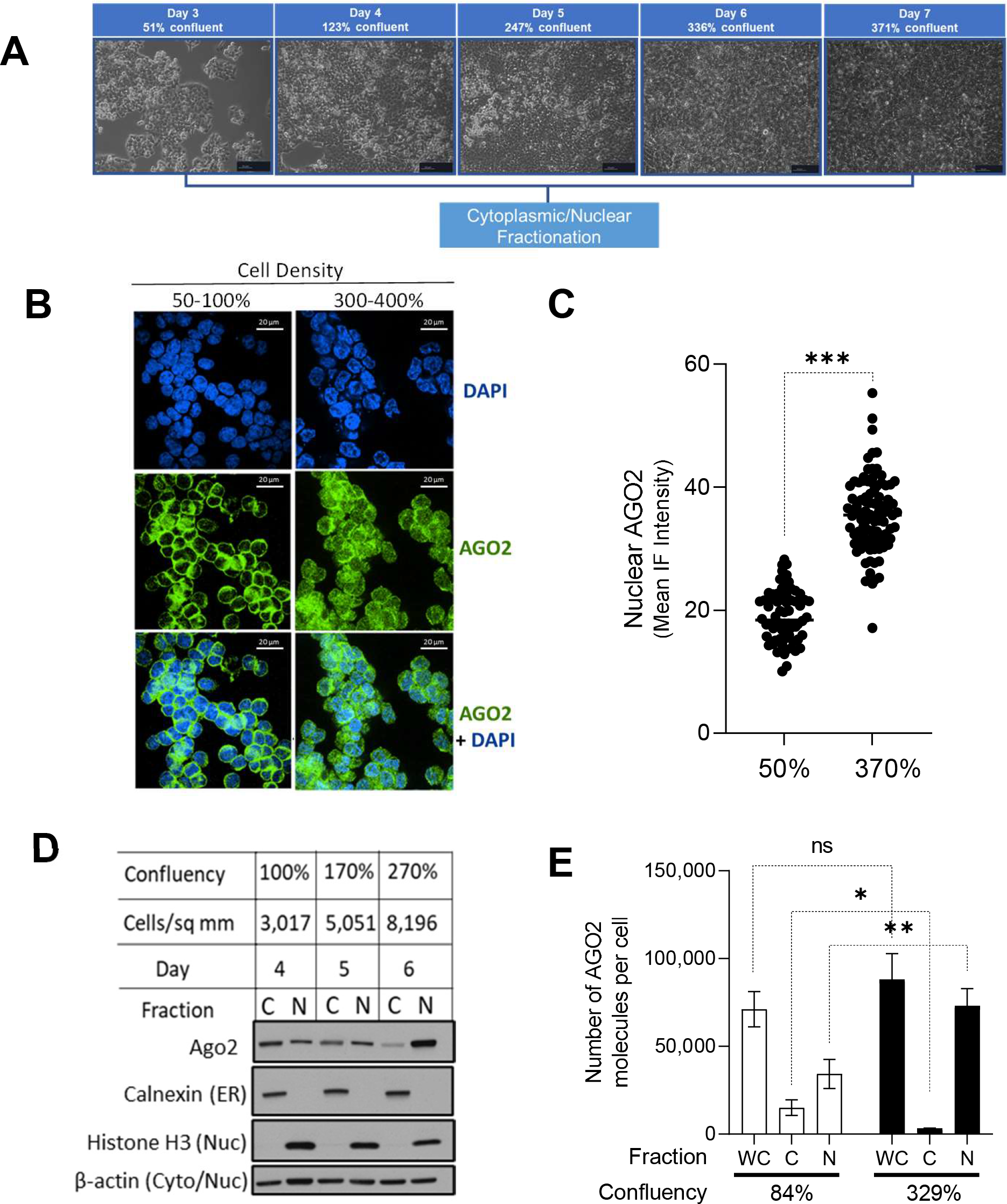
AGO2 shifts localization to the nucleus in cells grown to high density. (*A*) Microscopy images (0.25X magnification) taken immediately before harvesting each time point of increasing cell density stress. Average confluency calculated based on cell count and available surface area for growth. (B) Immunofluorescence of AGO2 (green) localization and DAPI staining of nuclei (blue) at low confluency (80-100%) and high confluency (300-400%). (C) Quantification of nuclear localization of AGO2 by mean immunofluorescence intensity. Unpaired t-test, two-tailed used to calculate p-value <0.0001. (D) Western blot showing AGO2 localization and purity of cytoplasmic (C) and nuclear (N) fractions (N=3). (*E*) Estimates of absolute copy per cell of AGO2 in whole cell (WC), cytoplasm (C), and nuclear (N) fractions determined by Western blot comparing densitometry of a serial dilution of recombinant AGO2 compared to signal from fractionated lysate. Unpaired t-test with Welch’s correction to calculate significance of the change in cytoplasm p=0.04 and nucleus p=0.006.

Cell viability can influence the integrity of cell membranes and nuclear membranes, so to ensure the shift in AGO2 localization was not due to cell death or ruptured membranes, we used dual viability staining with annexin V and propidium iodide followed by flow cytometry analysis to measure the percentage of viable, early apoptotic, and late apoptotic/dead cells. At the highest cell density on Day 7, over 90% of the cells were viable, suggesting that higher cell density and changing localization of AGO2 were not associated with increased apoptosis (**Supplementary Figure 1**). We observed similar altered localization of AGO1 (**Supplementary Figure 2**). The shift of AGO2 to cell nuclei at higher confluence was also observed in HEK293 and A549 cells where AGO2 is mostly cytoplasmic at normal confluence (**Supplementary Figure 3**). Increased nuclear localization was not observed in primary fibroblast GM02153 (apparently healthy individual), HeLa, or HepG2 cells.

To ensure the fractionation protocol was not introducing technical artifacts in the relative distribution, we then used absolute quantification to measure AGO2 protein in terms of numbers of proteins per cell as a function of increased cell density in whole cell, cytoplasm, and nuclei (**Figure 1E**). We used a serial dilution of recombinant AGO2 at known concentrations as a standard for titration (**Supplementary Figure 4**). This quantitative analysis confirmed the shift in cytoplasmic versus nuclear localization observed in our initial microscopy and western analysis (**Figure 1B-D**). Whereas the ratio of nuclear:cytoplasmic AGO2 was 2:1 when grown to 84% confluence, the ratio was 12:1 when grown to 329% confluence. The total amount of AGO2 (whole cell) inside cells at high and low confluence, was similar, ∼70,000 copies per cells. These data suggest that while higher cell density shifts AGO2 from cytoplasm to nucleus, overall expression levels do not significantly increase.

### Depletion of miRNAs in *DROSHA-/-* restricts AGO2 localization to the cytoplasm

DROSHA protein is responsible for the biogenesis of most miRNAs and knocking out *DROSHA* expression depletes most mature miRNAs (48,58). To evaluate how the decrease in miRNA expression affects localization of AGO2 as a function of cell density, we grew *DROSHA* knockout HCT116 cells at increasing confluence. In contrast to the nuclear enrichment observed in wild-type HCT116, AGO2 remained in the cytoplasm regardless of the level of confluence (**Figure 2A**). The overall amount of AGO protein decreases in DROSHA-/- cells, consistent with prior reports that AGO2 is unstable when not loaded with small RNAs (59-62).

**Figure 2.**
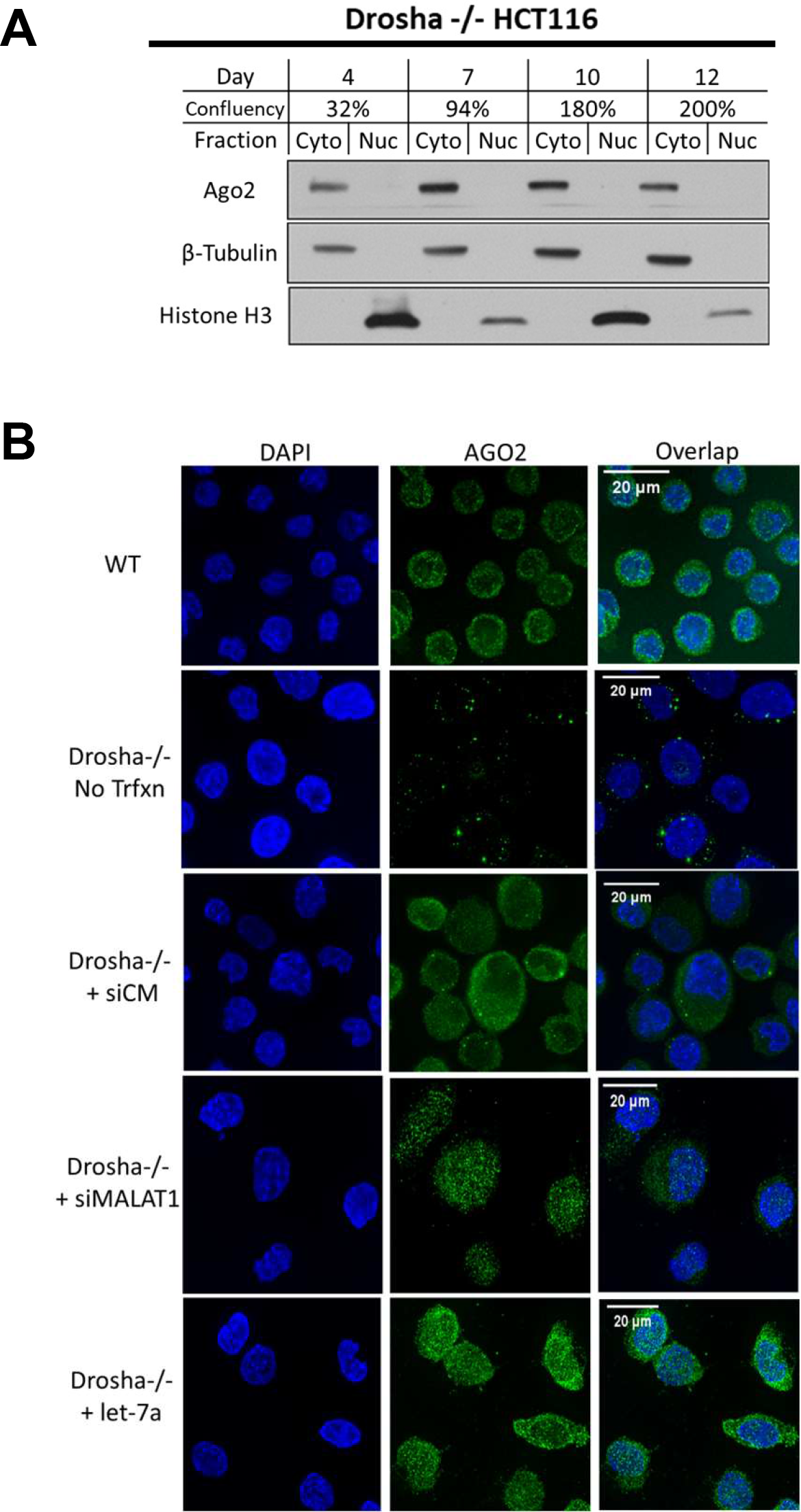
Depletion of miRNAs in DROSHA-/- restricts AGO2 localization to the cytoplasm. (A) Western blot showing AGO2 localization in isolated cytoplasm and nuclear fractions in Drosha-/- grown to high confluency (N=3). (B) Immunofluorescence microscopy showing AGO2 localization in WT HCT116, DROSHA-/- cells, and DROSHA-/- cells transfected with microRNA mimics and control oligos.

The restricted localization of AGO2 to the cytoplasm when miRNAs are depleted could be explained by at least two mechanisms. The first is that the loading of AGO2 with specific miRNAs are necessary for nuclear localization (63). The second is that unloaded AGO2 protein is unstable and degraded faster than nuclear localization (62). To discriminate between these hypotheses, we transfected small duplex RNAs into cells to determine their effect on the localization of AGO2. siMal-6 is a duplex siRNA known to silence the nuclear noncoding RNA MALAT-1 in wild-type cells, while let-7a is designed as a miRNA mimic.

We transfected duplex RNAs and observed increased nuclear localization relative to a negative control (lipid only) transfection (**Figure 2B**). These results are supported by western analysis (**Supplementary Figure 5**). Our observation that nuclear localization of AGO2 can be rescued when small RNA guides are available is consistent with previous findings suggesting that loading of small RNA into AGO2 occurs in the cytoplasm (10,11), and that loading is necessary for nuclear localization. Transfection of a negative control duplex RNA, siCM, that does not target a sequence found within the human genome also rescued AGO2 expression and nuclear localization (**Figure 2B, Supplementary Figure 5**). Taken together, these findings suggest that AGO2 requires a small RNA for nuclear localization and is likely unstable when unloaded.

### Loss of TNRC6A expression promotes AGO2 localization to the nucleus

Trinucleotide repeat containing protein 6 (TNRC6) is a family of three miRISC scaffolding proteins, TNRC6A, TNRC6B, and TNRC6C, that can bind up to three AGO proteins (64-66). TNRC6 is an essential protein partner for canonical miRNA activity by bridging interactions with miRISC accessory proteins affecting decapping, deadenylation, and mRNA decay (67,68). TNRC6 and AGO2 proteins are known to interact in both cytoplasm and nuclei and their interaction can influence subcellular localization (6,13,45,47,66,69-71). Although plant AGO1 proteins contain both a nuclear localization signal (NLS) and a nuclear export signal (NES) (49), metazoan AGO proteins lack a known NLS and NES. However, the metazoan-specific GW182 paralog, TNRC6A, was shown to possess both a functional NLS and NES (69).

Because of the important partnership between AGO2 and TNRC6 in mammalian cells, we examined how depleting the expression of TNRC6 would affect localization of AGO2 as a function of cell density. While we were not able to obtain a *TNRC6ABC* triple knockout cell line, the *TNRC6AB* double knockout cell line is viable, allowing us to use it to evaluate the impact of depleting most TNRC6 protein on the localization of AGO2. We observed an enrichment of AGO2 in the nucleus in TNRC6AB knockout cells regardless of cell confluence (**Figure 3A**). These data suggest that, while TNRC6 and AGO can form complexes in cell nuclei or in cell cytoplasm (13,47,56,66,69), full expression of TNRC6 is important for retaining AGO2 protein in the cytoplasm.

**Figure 3.**
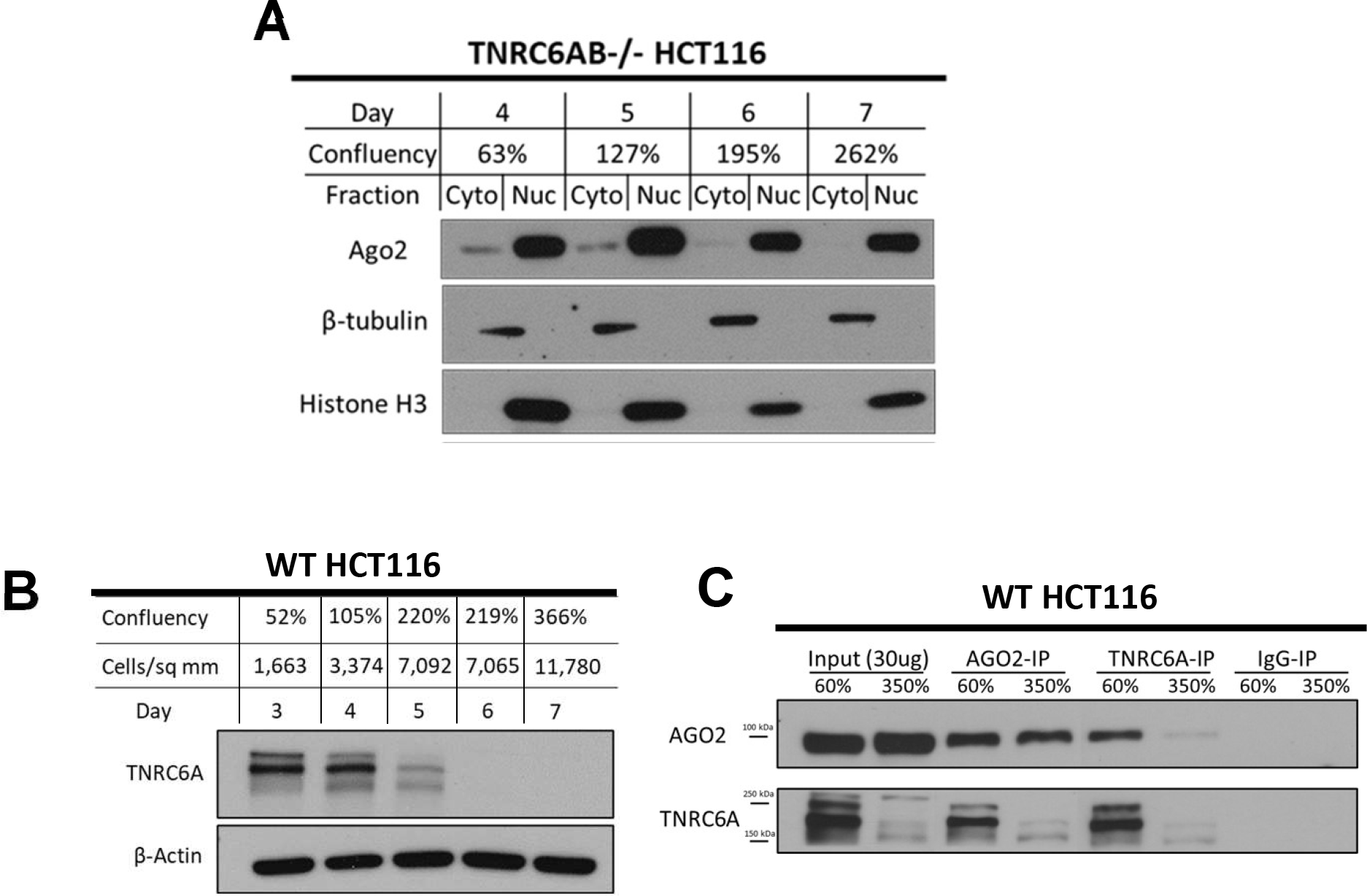
Loss of TNRC6A expression promotes AGO2 localization to the nucleus. (A) Western blot showing AGO2 localization in isolated cytoplasm and nuclear fractions in TNRC6AB-/- grown to high confluency (N=3). (B) Western blot showing TNRC6A in whole cell lysate in WT HCT116 cells grown to high confluency (N=3). (C) Immunoprecipitation of AGO2 and TNRC6A in WT HCT116 whole cell lysate at normal confluency (60%, Day 3) and high confluency (350%, Day 7) (N=3).

After establishing that the depletion of TNRC6 could affect AGO2 localization, we evaluated the effect of high cell density on expression of TNRC6A. TNRC6A was the focus of these experiments because it was the only TNRC6 paralog with known NES and NLS and for which we possessed an anti-TNRC6 antibody that was adequate for western analysis or immunoprecipitations. Western analysis revealed that growing HCT116 cells to high density caused a >90% decrease in TNRC6 protein expression (**Figure 3BC**).

To further test the impact of cell density on the AGO2:TNRC6 partnership, we used reciprocal co-immunoprecipitations with anti-AGO2 or anti-TNRC6A antibodies (**Figure 3C**). The input control samples reinforce the conclusion that TNRC6A protein expression decreases at high cell density. At normal cell densities, an anti-AGO2 antibody pulls down TNRC6A protein and an anti-TNRC6A antibody pulls down AGO2. At higher cell densities, however, an anti-AGO2 antibody pulls down less TNRC6 and the anti-TNRC6 antibody pulls down less AGO2. These experiments reveal decreased TNRC6A expression and a decreased potential for AGO2:TNRC6A interaction suggesting that high cell density perturbs formation of complexes that are responsible for canonical miRNA regulation. These findings support a hypothesis where TNRC6 proteins may serve as anchors retaining AGO in the cytoplasm, and the loss of TNRC6 expression by (1) CRISPR/Cas9-mediated knockout (**Figure 3A**) or (2) downregulation at high cell density (**Figure 3BC**) may contribute to nuclear localization of AGO.

### Global Increase in miRNA Expression at High Cell Density

We used small-RNAseq to evaluate the effect of high cell density on miRNA expression. The overall number of miRNA reads increased by ∼1.5 fold (**Figure 4A**). Consistent with the overall increase, when the expression of individual miRNAs was examined, most show a 1.5-2 fold increase (**Figure 4B**), and 80% of all miRNAs that show significant changes are up-regulated (**Supplementary Figure 6A**). The increase in miRNA expression agrees with previous reports that cell-cell contact or cell density activates miRNA expression (72-74).

**Figure 4.**
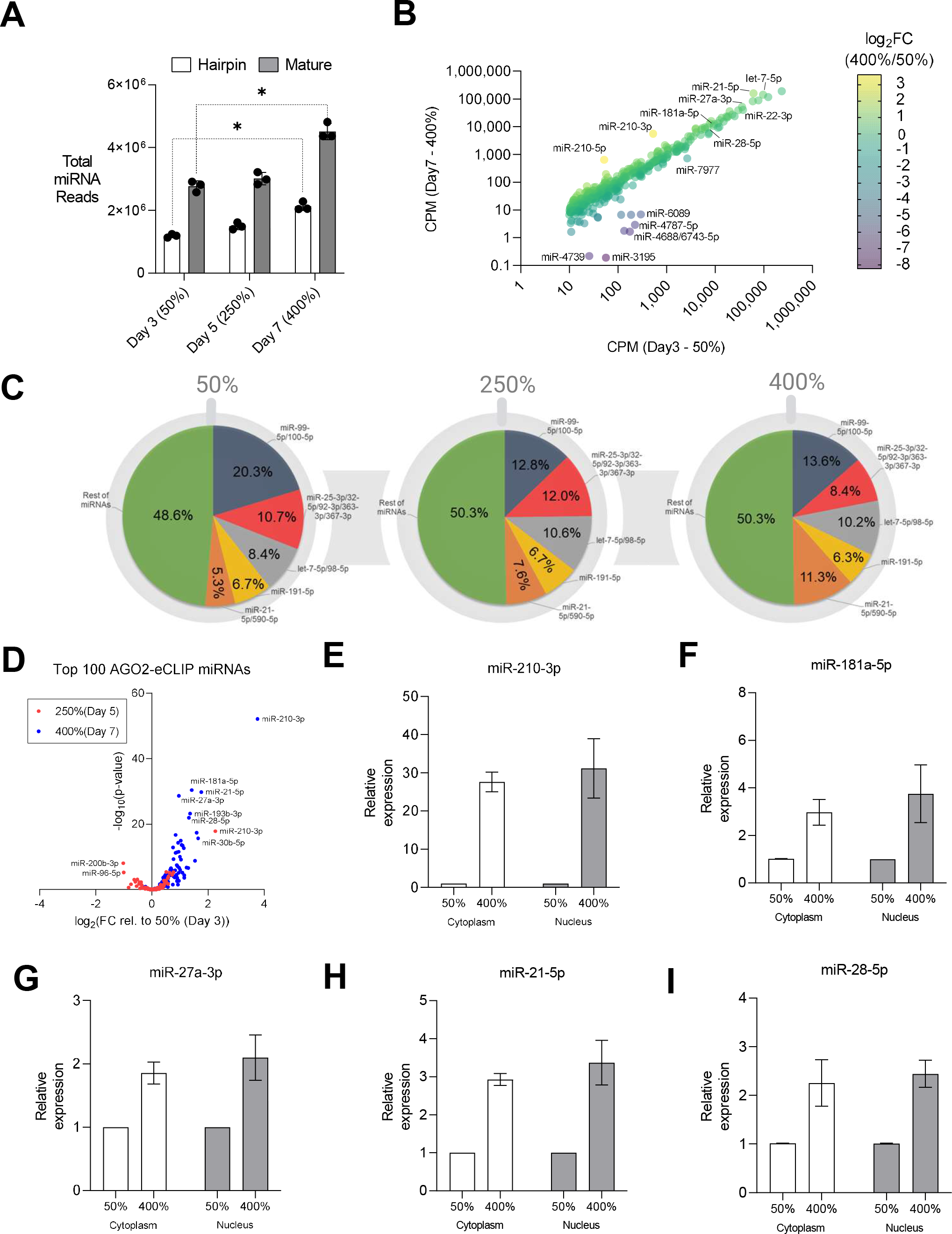
Global Increase in miRNA Expression at High Cell Density. (A) Global miRNA expression from small-RNA-seq (CPM). Paired t-test used to calculate significance with hairpin reads on day 7 relative to day 3 as p=0.012 and mature reads on day 7 relative to day 3 as p=0.019. (B) Total miRNA reads aligning to hairpin sequences and mature sequences. (C) miRNA expression from small-RNA-seq based on families sharing seed sequence at 50%, 250%, and 400% confluency. (D) Volcano plot showing significance and fold-change of the expression of the top 100 AGO2-eCLIP miRNAs at 250% confluency relative to 50% confluency (light blue) and at 400% confluency relative to 50% confluency (dark blue). (E-I) miRNA with greatest expression change in (D) measured by RT-qPCR in RNA isolated from cytoplasm and nuclear fractions at 50% and 400% confluency.

We observe that miRNAs belonging to five miRNA seed sequence “families” account for ∼50% of all sequencing reads in HCT116 cells regardless of confluence (**Figure 4C**). These data suggest that high cell density does not produce large changes in the relative regulatory potential of well-expressed miRNAs. Of the 100 miRNAs most closely associated with AGO2 from advanced immunoprecipitation experiments using the AGO2-eCLIP-seq method (33,48), we found that most showed increased expression (**Figure 4D**). To validate this small-RNAseq data, we selected five abundant miRNAs associated with AGO2 from AGO2-eCLIP-seq, miR-210-3p, miR-181a-5p, miR-21-5p, miR-27a-3p, miR-28-5p, that had the greatest increase in expression at high cell density relative to low cell density for analysis by qPCR (**Figure 4D**).

miRNA-210-3p, which has previously been reported to be up-regulated in hypoxic conditions (75-79), showed the largest increase (30-fold) in expression of any highly expressed miRNA (**Figure 4D**). All five miRNAs that were measured by qPCR showed increases in relative expression that were consistent with those observed by small-RNAseq (**Figure 4E-I**).

Since AGO2 was enriched in the nucleus at high cell density, we hypothesized that specific miRNAs that show the greatest increase in expression at high cell density might also become enriched in the nucleus. For all miRNAs examined, the increase in expression was the same regardless of whether expression was measured in nuclei or cytoplasm (**Figure 4E-I**). These qPCR data support the findings from our small-RNAseq analysis that changes in the expression of most, but not all, individual miRNAs at high cell density are modest. Interestingly, the 1.5-2-fold increase in global miRNA expression (**Figure 4A**) is similar to the increase in whole cell AGO2 expression (**Figure 1E**) (**Supplementary Figures 4 and 6BC**). The similar modest increases in AGO2 and miRNA expression suggests that some mature miRNAs may be stabilized by loading into AGO for miRISC complex formation, and this is supportive of previous work investigating global miRNA activation at high cell density (72).

### Nuclear localization of AGO2 relieves repression of cytoplasmic 3’UTR targets

The shift of AGO2 protein to the nucleus in response to changes in cell density might affect canonical miRNA repression that relies on cytoplasmic AGO protein. To test this hypothesis, we examined the impact on target genes that were candidates for canonical regulation through binding of miRISC within their 3’-untranslated regions (3’-UTRs) (**Figure 5A**). To identify candidate targets of miRISC regulation, we used a multi-pronged transcriptomics approach that first incorporated enhanced crosslinking immunoprecipitation sequencing (eCLIP-seq) using an anti-AGO2 antibody to identify candidate target mRNAs that physically associate with AGO2 in the cytoplasmic fraction within the 3’UTR (33,80).

**Figure 5.**
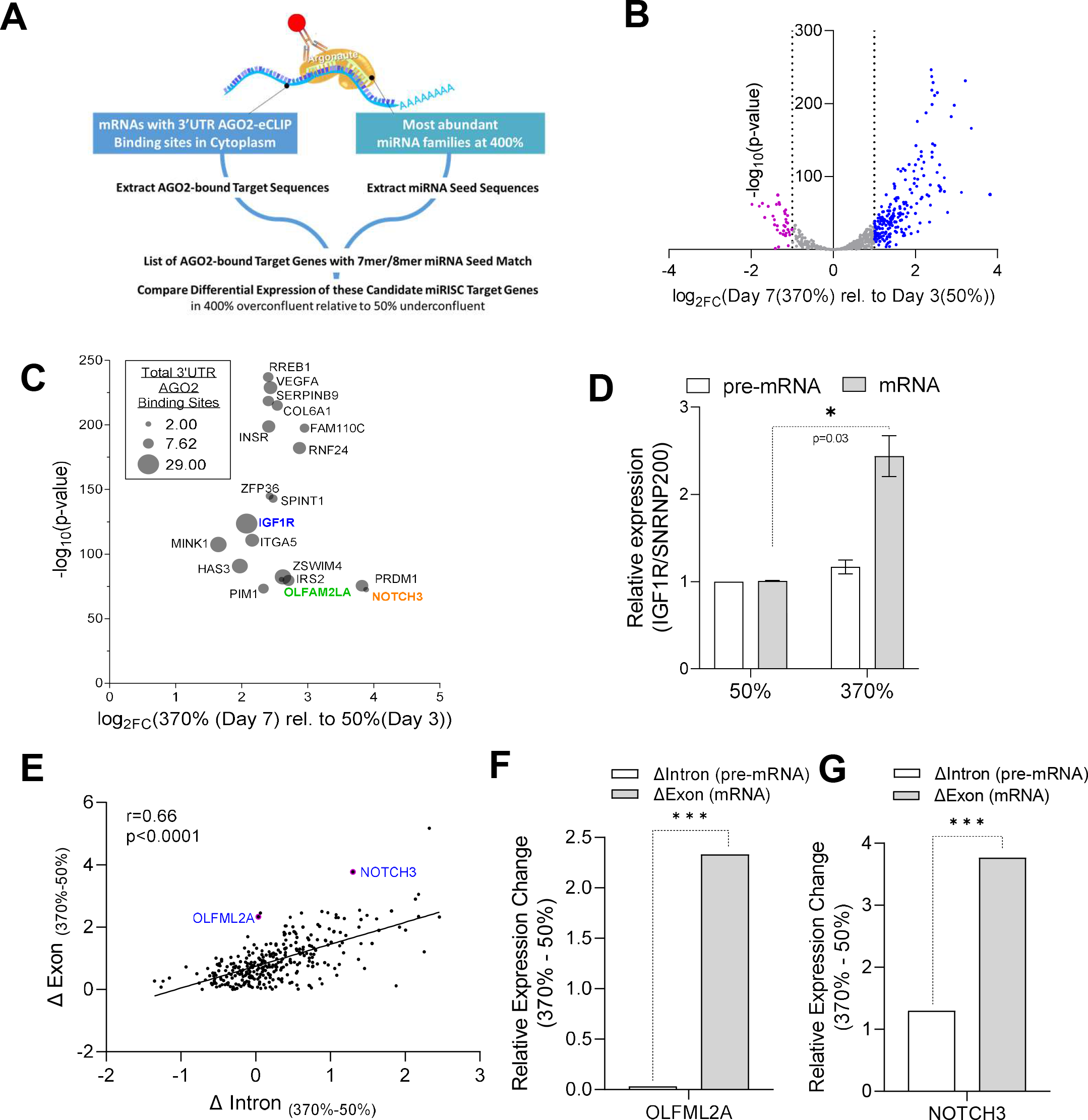
Nuclear localization of AGO2 relieves repression of cytoplasmic 3’UTR targets. (A) Candidate refinement using AGO2-eCLIP sequencing, small-RNA-sequencing, miRNA family seed alignment, and whole transcriptome RNA-sequencing. (B) Volcano plot of differential expression from RNA-seq of candidates with abundant miRNA family seed matches to 3’UTR AGO2-eCLIP binding sites in the cytoplasm. (C) Plot showing differential expression, significance, and number of cytoplasmic AGO2-eCLIP binding sites within 3’UTR of candidate genes targeted by miRISC. (D) Basal expression level of pre-mRNA and mRNA for IGF1R measured with RT-qPCR from RNA harvested on Day 3 (underconfluent, 50%) and Day 7 (overconfluent, 370%). Significance denoted with *, p-value equals 0.03. (E) Exon-Intron Split Analysis (EISA) of AGO2-eCLIP candidates with sites of 3’UTR occupancy and abundant miRNA family seed matches. (F) EISA analysis showing changes in expression of intronic reads (pre-mRNA) and exonic reads (mRNA) at high density (370%) relative to low density (50%) for (F) OLFML2A, p-value = 3.32 × 10^-11, FDR=1.47 × 10^-8 and (G) NOTCH3, p-value = 2.75 × 10^-11, FDR = 1.37 × 10-8.

The next criteria required that these regions of AGO2 occupancy from AGO2- eCLIP on candidate target mRNAs have strong, 7-mer or 8-mer, complementarity to seed sequences belonging to the top 20 most abundant miRNA families from small-RNAseq at high cell density (**Fig. 4CD, 5A, Supplementary Figure 7**). Finally, we used whole transcriptome RNAseq (RNAseq) (**Supplementary Figures 8 and 9**) to examine the differential expression of these candidate miRISC targets at high cell density (400%) relative to low cell density (50%). In contrast to global expression changes, we found that the subset of candidate target genes that possess binding sites for miRISC showed increased expression at high cell density (**Fig. 5B, Supplemenary Figure 9BC**). This outcome is consistent with the hypothesis that relocation of AGO protein to the nucleus reduces canonical repression by miRNAs in the cytoplasm.

We hypothesized that miRISC target genes undergoing canonical miRNA regulation should have the greatest effect at the post-transcriptional level. Candidate miRISC target genes that possessed several AGO2-eCLIP binding sites aligning with strong miRNA seed complementarity and had a significant increase in expression at high versus low cell density were selected for further analysis (**Fig. 5C**). One of the top candidate targets, *IGF1R*, possessed twenty-nine AGO2-eCLIP binding sites within its 3’UTR in the cytoplasm, more than any other gene. Quantitative PCR (qPCR) revealed upregulation of mature *IGF1R* mRNA, but not pre-mRNA (**Fig. 5D**), consistent with de-repression occurring at the post-transcriptional level and consistent with possible reversal of canonical miRNA-mediated repression.

To separate transcriptional effects from post-transcriptional effects for multiple potential miRISC targets across the transcriptome, we used Exon-Intron Split Analysis (EISA) (53) to investigate all the candidate targets with increased expression at high cell density (blue dots in **Figure 5B**). The EISA analysis uses whole transcriptome RNAseq data to separate the transcriptional and post-transcriptional components of gene regulation by comparing the amounts of exonic and intronic reads from expressed mRNA transcripts. Changes in intronic read counts or pre-mRNA reads predict changes in upstream transcriptional activity, and changes in downstream post-transcriptional regulation can be predicted from the presence of changes in exonic read counts or mature mRNA reads and the absence of changes in intronic read counts (53).

We hypothesized that if the shift of AGO2 to the nucleus at high cell density results in de-repression of these cytoplasmic targets, we would observe an increase in the mature mRNA levels indicating a post-transcriptional effect. These target genes would have the greatest increase in exonic reads (reads aligning to changes in mature mRNA along the y-axis) and have the least increase in intronic reads (reads aligning to changes in pre-mRNA along the x-axis). The candidate genes that fit these criteria were found plotted above the best fit line, indicating that these candidate miRISC target genes showed a greater increase in gene expression at the post-transcriptional level than at the transcriptional level (**Figure 5E, Supplementary Figure 10**). EISA cannot be used with genes that overlap, precluding measurement at the *IGF1R* locus and at some of the other candidate miRISC targets identified in **Figure 5BC**. EISA analysis revealed several candidate genes were upregulated primarily at the post-transcriptional level, with the greatest increase in exonic reads along the y-axis observed for *OLFM2LA* (p-value=3.32 × 10^-11^, FDR=1.47 × 10^-8^) and *NOTCH3* (p-value=2.75 × 10^-11^, FDR=1.37 × 10^-8^) (**Figure 5E-G**). Several other upregulated candidate miRISC targets showed a greater increase in intronic reads along the x-axis suggesting upregulation arising from upstream transcriptional effects (**Figure 5E, Supplementary Figure 10ABCD**), and these genes may be worth investigating further in the future to separate potential direct and indirect effects of AGO2 nuclear localization on the regulation of transcription.

### miRISC target genes have longer half-lives in the cytoplasm when AGO2 is shifted to the nucleus

Inhibition of gene expression by miRNAs is caused by repressing translation or by reducing the stability of the target RNA, with the latter mechanism being dominant (81-83). Because reduced RNA stability is a hallmark of miRISC action, we evaluated the stability of candidate mRNA targets at different cell densities in cytoplasm and nuclear fractions (**Figure 6A**). We chose to examine *IGF1R* because it possessed the highest number of potential binding sites for miRISC complexes and showed an increase in expression for mature mRNA (**Supplementary Figure 7 and Figure 5CD**). *NOTCH3* and *OLFMLA2* were examined because they had the greatest increase in mature mRNA expression revealed by EISA (**Figure 5E-G**).

**Figure 6.**
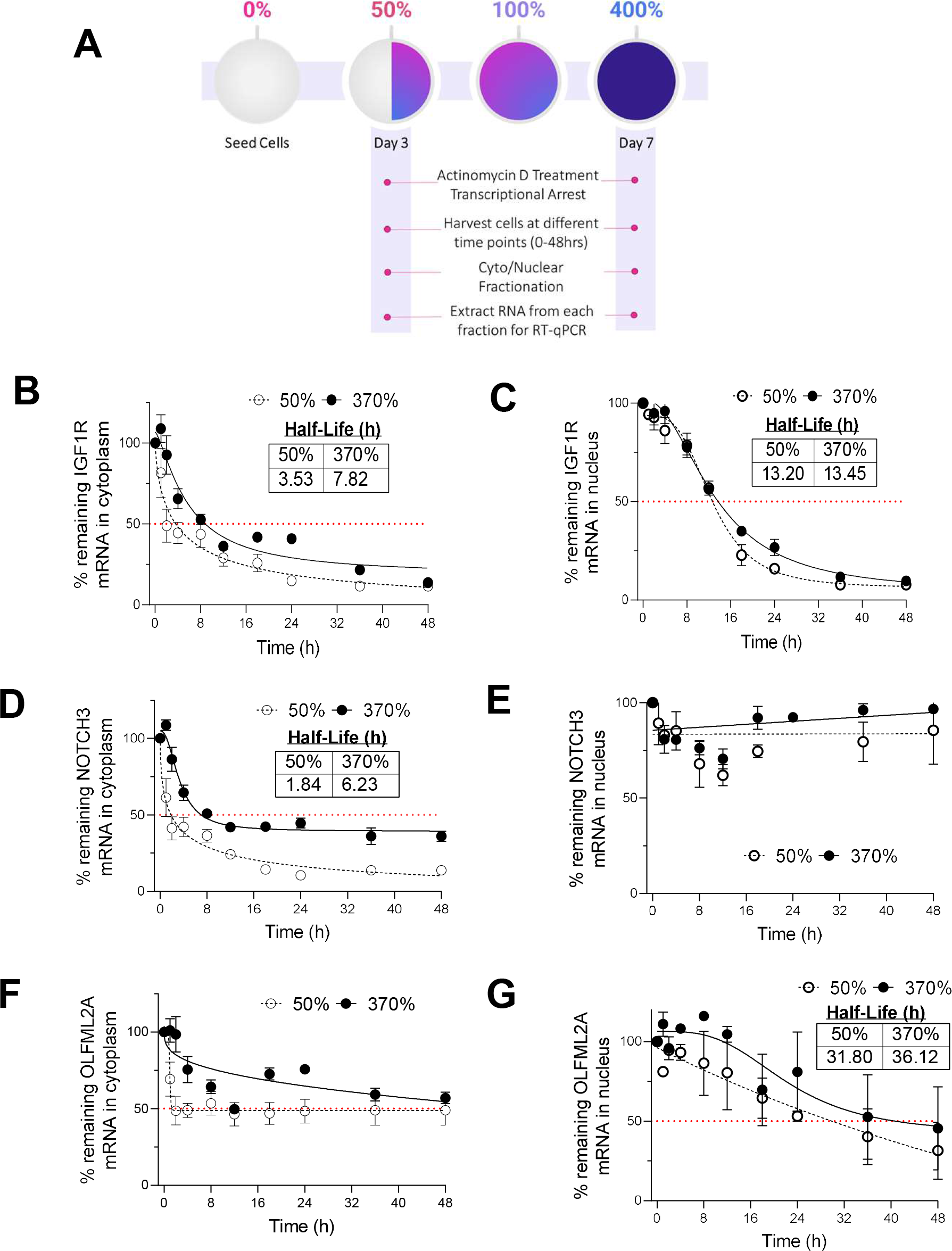
miRISC target genes have longer half-lives in the cytoplasm when AGO2 is shifted to the nucleus. (A) Scheme for measuring RNA stability at low confluency (50%) and at high confluency (400%) in RNA isolated from cytoplasm and nuclear fractions at each time point (0-48 hrs). (B-E) mRNA stability assays for upregulated miRISC targets in RNA isolated from cytoplasm (B, D, F) and in RNA isolated from nuclei (C, E, G).

To measure the effect of AGO2 localization on the stability of these target mRNAs, we added actinomycin D (actD) to arrest transcription in cells that were grown to 50% and 370% confluency, harvested the cells at several time points over forty-eight hours after actD treatment, fractionated the cells at each time point, and extracted RNA from each fraction at each time point for analysis by RT-qPCR (**Figure 6A**). We validated that the actD treatment arrested pre-mRNA transcription at the same time for cells grown to 50% or 370% confluency (**Supplementary Figure 11A-C**). These RNA stability assays revealed that *IGF1R, NOTCH3, and OLFM2LA* mRNAs had longer half-lives in the cytoplasm at high cell density than at lower confluence (**Figure 6BDF**), consistent with loss of miRNA-mediated mRNA decay in the cytoplasm when cells are at high confluence and AGO2 is shifted to the nucleus. By contrast, there was no significant change in the stability of these mRNAs in the nucleus (**Figure 6CEG**). Some candidate miRISC targets that were evaluated did not show a change in mRNA stability or had too long or too short half-lives to be measured (**Supplementary Figure 12**).

### Constitutive nuclear localization of AGO2 (NLS-AGO2) phenocopies relieved miRNA repression

It is possible that growing cells to extreme, high-density conditions can trigger changes to gene expression that are independent of AGO2 localization and miRNA-mediated regulation. To determine if the upregulation or potential de-repression of the candidate miRISC targets is directly related to nuclear localization of AGO2, we obtained CRISPR/Cas9-engineered cells that express AGO2 protein with a nuclear localization sequence (NLS) added to the N-terminus. These were knock-in cells with NLS added to both alleles of the endogenous AGO2 gene (NLS-AGO2 HCT116) (**Supplementary Figure 13**), avoiding overexpression and disruption of endogenous stoichiometry while genetically forcing nuclear localization of AGO2. Mammalian AGO proteins lack a known, endogenous NLS sequence, so we used the SV40 large T-antigen NLS sequence that was previously reported to manipulate AGO2 localization (13,47) (**Supplementary Figure 13**).

AGO2 protein is localized to the nucleus in the NLS-AGO2 cell line (**Figure 7AB**). We examined mRNA expression of three candidate miRISC target genes, *IGF1R*, *NOTCH3*, and *OLFML2A* (**Figures 5,6**). All three candidate targets mRNAs showed increased expression in NLS-AGO2 engineered cells relative to wild-type cells (**Figure 7C-E**). This result is consistent with the conclusion that relocalization of AGO2 to the nucleus reduces the cytoplasmic pool of AGO protein and reduces miRNA-mediated repression of these candidate targets.

**Figure 7.**
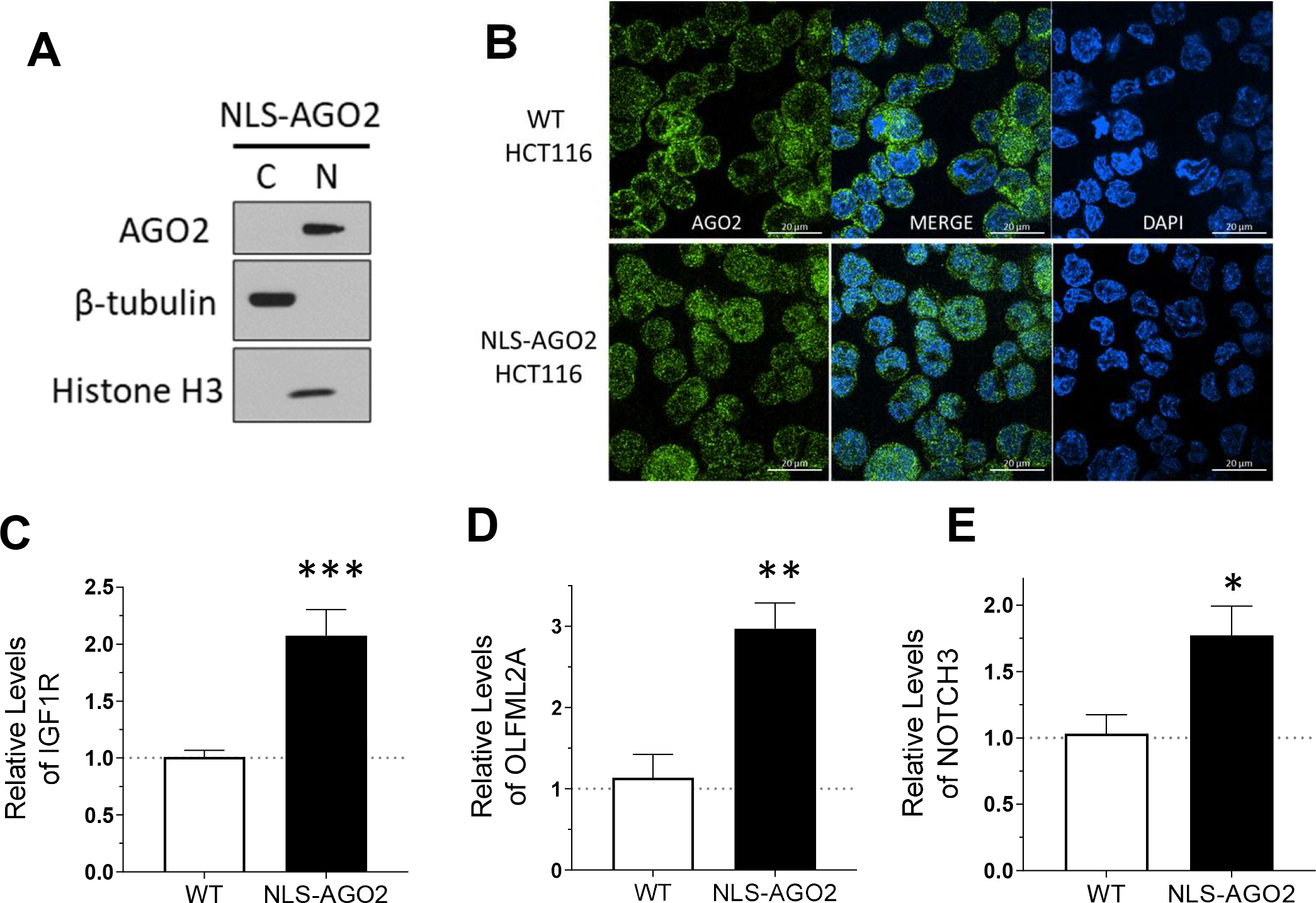
Constitutive nuclear localization of AGO2 (NLS-AGO2 knock-in) phenocopies overconfluent culture. (A) Western blot validation of AGO2 nuclear localization in CRISPR/Cas9-NLS-AGO2 knock-in HCT116 (NLS-AGO2). (B) Immunofluorescence validation of AGO2 nuclear localization in NLS-AGO2 compared to WT HCT116. (C, D, E) RT-qPCR results describing top candidate miRISC target expression changes in NLS-AGO2 cell line relative to WT (N=3). Significance denoted by *’s with (C) IGF1R, p=0.0004, (D) OLFML2A, p=0.005, (E) NOTCH3, p=0.03.

### Constitutive cytoplasmic localization of AGO2 (NES-AGO2) is sufficient to rescue repression

In contrast to the upregulation or potential de-repression observed with nuclear localization of AGO2, we hypothesized that restoring AGO2 localization in the cytoplasm should be sufficient to repress the expression of the candidate miRISC targets. To test this hypothesis, we evaluated the effects of directing AGO2 to the cytoplasm in cells grown to normal confluence using AGO1/2/3 knockout cells transfected with either a plasmid encoding wild-type AGO2 or AGO2 with a nuclear export sequence found in plant AGO1 (49) added to its N-terminus (NES-AGO2) (**Figure 8A**).

**Figure 8.**
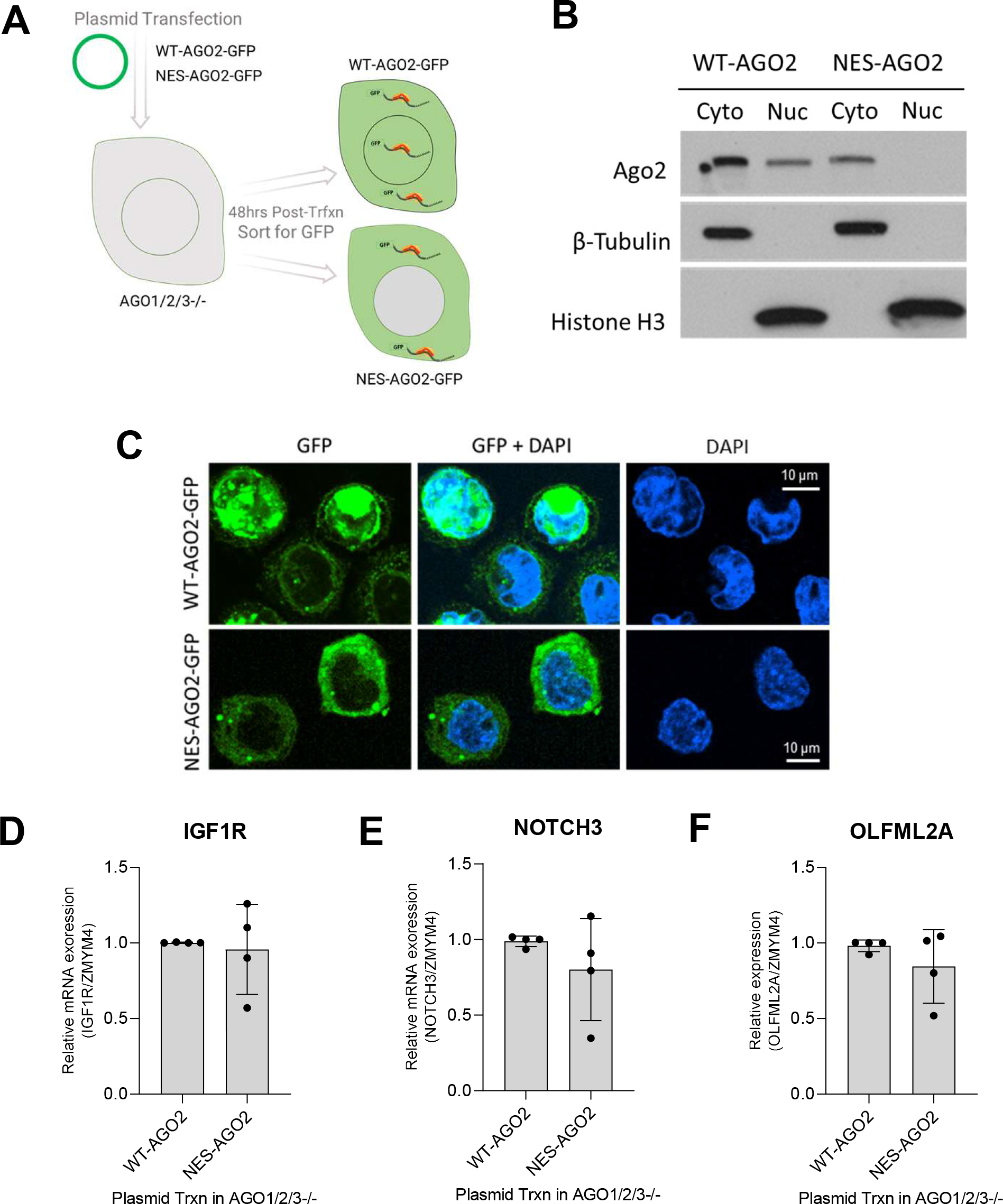
Constitutive cytoplasmic localization of AGO2 sufficient to rescue repression. (A) Scheme showing transfection of WT-AGO2-GFP and NES-AGO2-GFP in AGO1/2/3-/- cells. (B,C) Expression and localization of WT-AGO2-GFP and NES-AGO2 evaluated by western or microscopy. (D-F) Expression of WT-AGO2-GFP and NES-AGO2 restores repressed expression of IGF1R, NOTCH3, and OLFML2A.

To isolate the cells that successfully restored AGO2 expression, the plasmids also contained a GFP tag on the N-terminus of AGO2 allowing transfected cells to be sorted with flow cytometry and analyzed. Cells transfected with the NES-AGO2 plasmid were compared to cells transfected with plasmid expressing wild-type AGO2 (WT-AGO2), and both complementation experiments showed that AGO2 predominantly localized to the cytoplasm (**Figure 8BC**). In contrast to the NLS-AGO2 cells, where expression of *IGF1R*, *NOTCH3*, and *OLFM2LA* were increased, mRNA expression of these candidate miRISC targets were similar in the cells expressing WT-AGO2 or NES-AGO2 (**Figure 8D-F**). These data, from cells that express WT-AGO2, NLS-AGO2, and NES-AGO2, support the hypothesis that the ability for AGO2 to repress the candidate miRISC targets is dependent on the amount of AGO2 protein present in the cytoplasm.

### Shift of AGO2 to the nucleus is associated with dynamic changes in pathways and gene networks related to cell adhesion, signaling, and migration

miRNAs can directly affect a specific phenotype, and many different miRNAs can also work together to collectively target a larger network of different genes within common pathways through weak but pervasive interactions (84,85). We used our RNAseq data for pathway analysis to evaluate how cellular processes change when AGO localizes to cell nuclei. This analysis revealed that candidate miRISC targets (**Figure 5AB**) that possess AGO2 binding sites as determined by our eCLIP data were enriched in pathways involved in cell signaling, communication, and migration (**Figure 9A**). Almost all these pathways are associated with changes in *IGF1R* expression, and some are also associated with changes in *NOTCH3* and *OLFML2A* levels.

**Figure 9.**
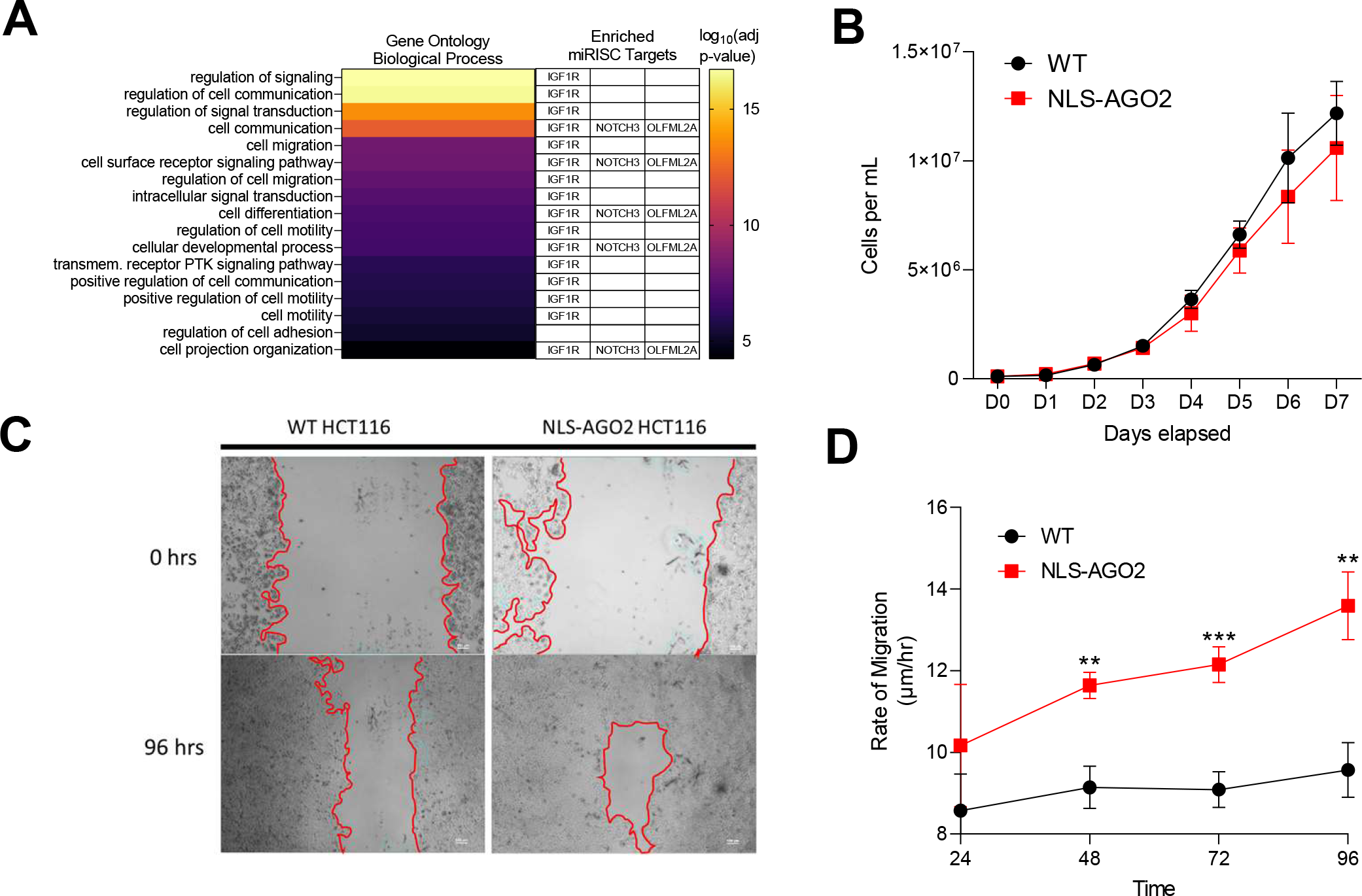
Shift of AGO2 to the nucleus is associated with dynamic changes in pathways and gene networks related to cell adhesion, signaling, and migration. (A) Pathway analysis of all upregulated cytoplasmic gene candidates with 3’UTR AGO2-eCLIP binding sites. Pathways related to IGF1R, NOTCH3, and OLFM2LA are shown in table. (B) Proliferation assay to measure cell growth rates of WT HCT116 compared to NLS-AGO2 HCT116. No significant difference. (C) Microscopy images from Scratch/Migration Assay showing movement of WT and NLS-AGO2 HCT116 cells grown for 96 hours. (D) Migration rate of WT and NLS-AGO2 HCT116 (N=3). Unpaired t-test was used to calculate significance. No significance at 24 hrs, 48 hrs: p=0.002, 72 hrs: p=0.001, 96 hrs: p=0.002.

To identify which networks of genes are affected by the loss of miRNA regulation in the cytoplasm, we performed gene network analysis. Gene network analysis also revealed that the differential expression of several candidate miRISC targets were positively correlated with one module of related pathways (Module: darkseagreen4, Pearson correlation 0.83, p<0.01, **Supplementary Figure 14A**) This module was enriched with genes related to cell adhesion, cell-cell junction assembly, intracellular signal transduction, and positive regulation of migration (p< 0.0001, **Supplementary Figure 14B**).

To begin investigating the potential impact of nuclear enrichment of AGO2 on cell migration, we used a scratch assay to compare wild-type and NLS-AGO2 HCT116 cells. The scratch assay is a simple cell culture procedure in which a thin wound or scratch is made on the surface of a dish followed by monitoring regrowth of cells over time. Wild-type and NLS-AGO2 cells grow at similar rates under normal cell culture conditions (**Figure 9B**). After the scratch is introduced (**Figure 9C**), however, the NLS-AGO2 cells showed increased migration relative to wild-type HCT116 cells (**Figure 9CD**). This result is consistent with the pathway analysis and gene network analysis that suggested redirecting the AGO2 component of the cellular pool of AGO protein to the nucleus can affect the expression of genes involved in cell migration.

### AGO2 is Enriched in the Nucleus in 3-D Tumor Spheroid Culture and Tumor Tissue

Growing cells to high density was a simple model system to mimic conditions found within a solid tumor. However, we were interested in more clinically relevant models to bridge the gap toward *in vivo* validation. Three-dimensional culture of tumor-derived cells is a better system to represent extracellular matrix organization, cell to cell interactions, and ability to form adhesion structures (86). We grew HCT116 cells in a collagen matrix to form tumor spheroids (**Figure 10A**) to test whether the nuclear localization of AGO2 would be affected in culture conditions that better represent characteristics of human tumors. We fractionated nuclei and cytoplasm and observed almost complete localization of AGO2 to cell nuclei (**Figure 10B**). RNA levels of potential target genes *IGF1R*, *NOTCH3*, and *OLFML2A* were increased in 3-D tumor spheroid culture (**Figure 10C-E**) compared to cells grown in 2-D culture to normal, ∼70% confluence. The increase or potential de-repression of the target mRNAs in 3-D tumor spheroids phenocopied the increase observed for cells grown to high density (**Figure 5C**) and NLS-AGO2 cells that had been engineered to have AGO2 localized to cell nuclei rather than cytoplasm (**Figure 7A**).

**Figure 10.**
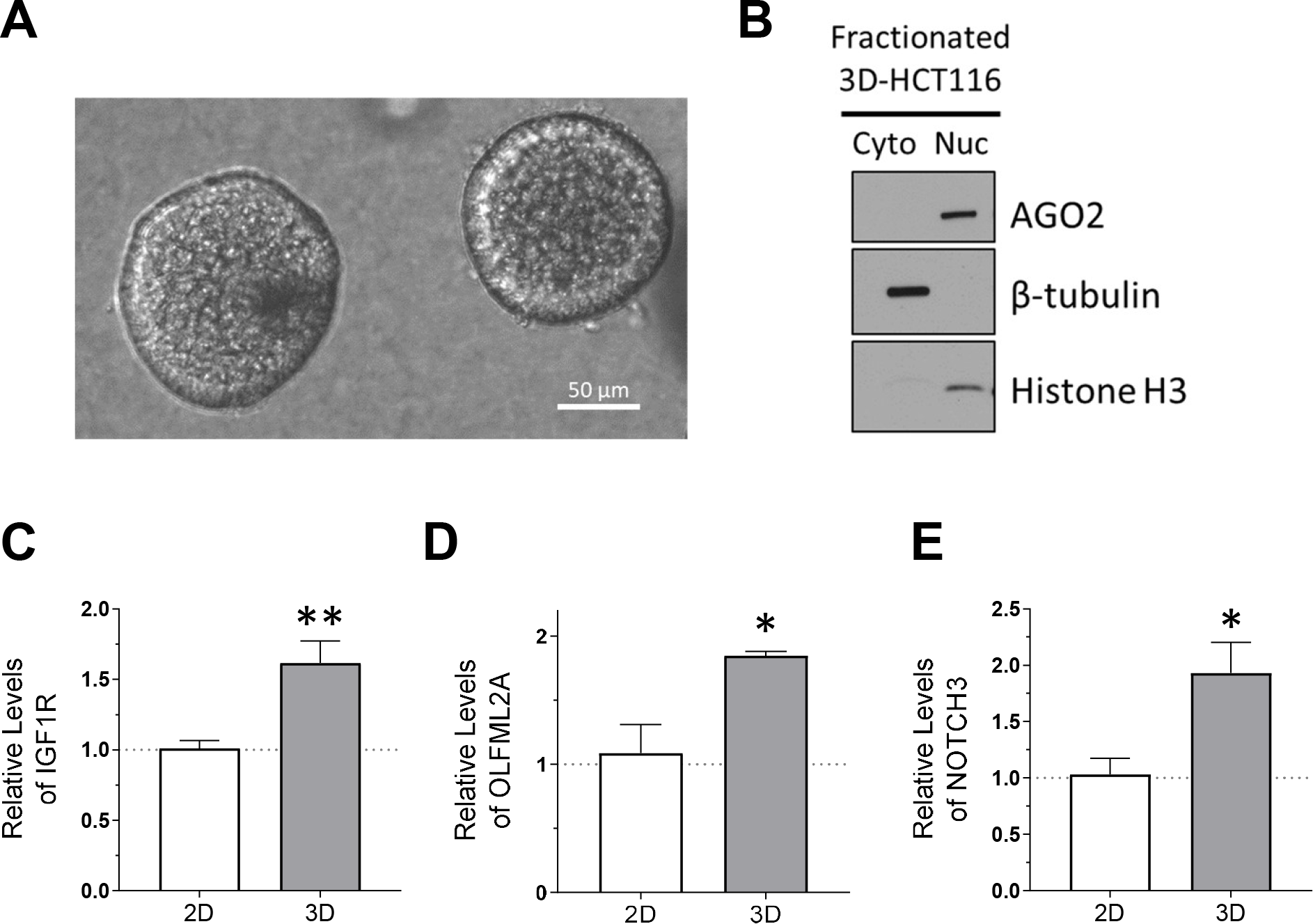
AGO2 is enriched in the nucleus in 3D-cultured tumor spheroids. (A) Microscopy images of 3D-cultured HCT116 tumor spheroids. (B) Western blot showing AGO2 localization in 3D-HCT116 (N=3). (C-E) Relative mRNA expression of candidate miRISC targets in 3D cultured HCT116 relative to parental 2D-cultured HCT116 cell line used to seed 3D culture. Significance denoted with *’s: (C) IGF1R, p=0.0016, (D) OLFML2A, p=0.036, (E) NOTCH3, p= 0.025.

The observed nuclear localization of AGO2 in 3-D tumor spheroid culture, in combination with our observation of changes in the expression of miRISC target genes associated with cell migration, encouraged us to next examine AGO2 localization in human colon tumor tissue. Localization in colon cancer tumor tissue was compared to that in adjacent normal tissue from the same patient. Tissue was obtained from twenty-eight patients representing multiple grades of colon cancer (**Figure 11A, Supplementary Fig. 15**). Methods were refined to separate the small amounts of human tissue available into cytoplasmic, nuclear, and chromatin fractions. We performed western analysis to evaluate AGO2 localization.

**Figure 11.**
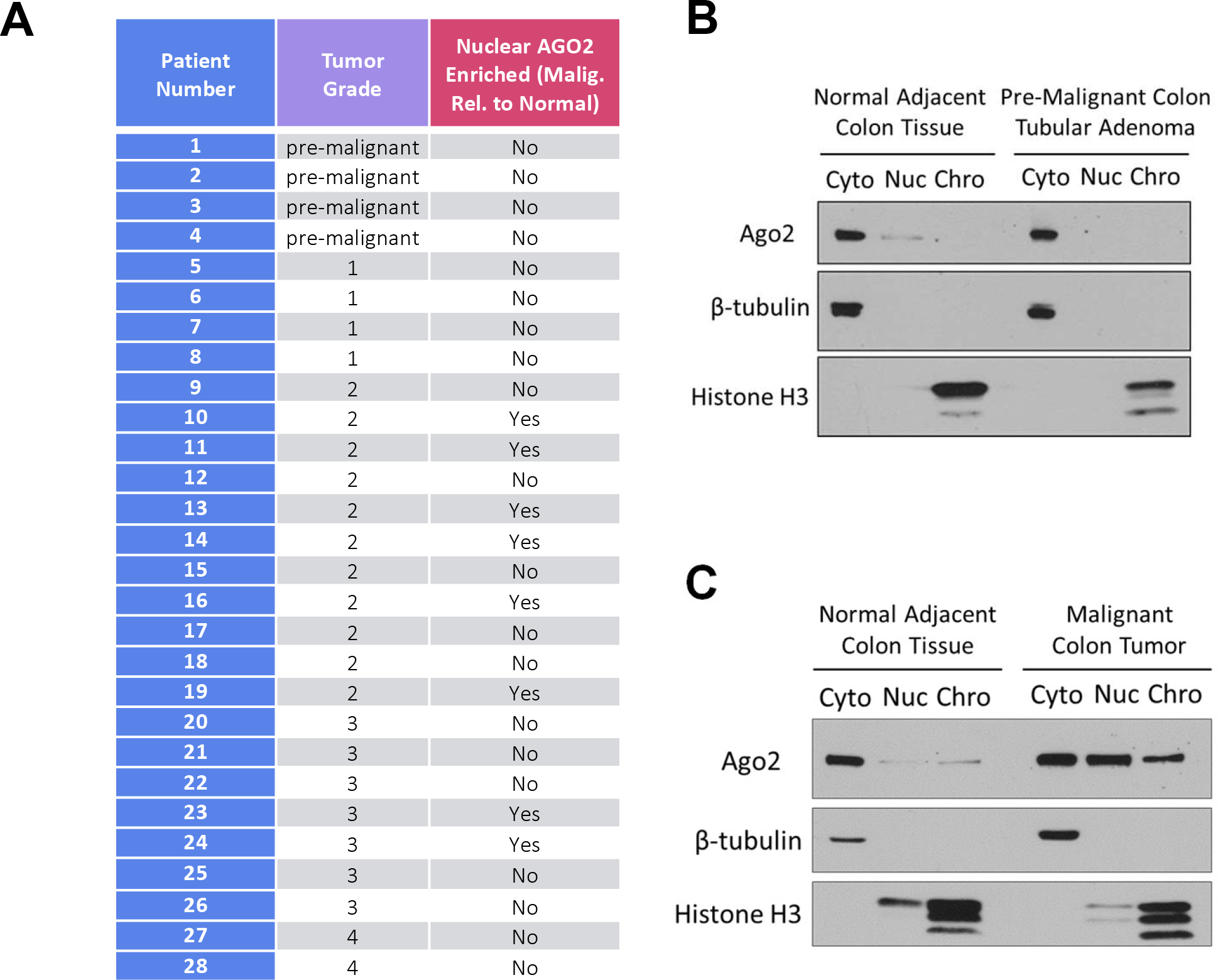
AGO2 localization in primary malignant, pre-malignant, and normal adjacent colon patient tissue. (A) Table describing 28 patients used to examine nuclear localization of AGO2. (B) Western blot showing AGO2 localization in cytoplasm (cyto), soluble nuclear (nuc), and chromatin (chro) fractions in normal colon tissue (left) and in pre-malignant colon adenoma tissue (right) from patient #2. (C) Western blot showing AGO2 localization in normal colon tissue (left) and in malignant colon adenocarcinoma tissue (right) from patient #10.

The tumor samples yielded different outcomes for the nuclear localization of AGO2. Of the twenty-eight samples, twenty-four had detectable nuclear AGO2 (**Supplementary Figure 15**). As part of our analysis, all samples were re-evaluated by hematoxylin and eosin staining (H&E staining) to ensure correct classification with regards to cancer grade and pathology. Eight samples showed enrichment of nuclear AGO2 in tumor tissue versus samples taken from normal adjacent colon tissue (**Figure 11A**). For example, based on pathology review of H&E-stained tissue sections (**Supplemental Figure 16**), sample 2 was determined to be from a pre-malignant, colon tubular adenoma that showed no nuclear AGO2 in nuclei nor was nuclear AGO detected in the matching normal adjacent tissue (**Figure 11B**). By contrast, sample 10 from a malignant, colon adenocarcinoma tumor tissue showed a distinct enrichment of nuclear AGO2 relative to normal colon tissue (**Figure 11C**). While the sample size is too small to draw definitive conclusions, all eight patient samples that were judged pre-malignant or grade one did not show nuclear enrichment, while 8/20 (40%) of samples from patients with grades 2-4 disease had nuclear-enriched AGO. These results suggest that nuclear localization of AGO2 can be enriched in some colon tumor samples, but that enrichment is not universal.

## DISCUSSION

### RNAi: cytoplasmic and nuclear

RNAi is a powerful natural regulatory mechanism for controlling post-transcriptional gene expression in the cytoplasm of various organisms (1,2). This cytoplasmic control of gene expression by duplex RNA has been used as an experimental tool (87) and a source of therapies (88). By contrast, potential nuclear functions of RNAi and even the presence of RISC factors in cell nuclei have been less obvious (89-92).

Improved purification protocols that exclude endoplasmic reticulum from nuclear lysate, however, demonstrated that AGO protein and miRNAs were present in cell nuclei (10,11,13,16,34,42,44,47,63,78,93). miRNAs were demonstrated to load in cell cytoplasm prior to transport into the nucleus. AGO can form a complex with TNRC6 in both the cytoplasm and the nucleus (13,11,28,66). TNRC6, a multidomain scaffolding protein, can then form complexes with proteins involved in mRNA decay, transcription, splicing, and other nuclear processes (13,66). Synthetic RNAs have the potential to control gene transcription or splicing (32,94). While these data demonstrate that synthetic RNAs can control nuclear gene expression and that RISC factors exist in cell nuclei, the potential for RNAi to exploit these mechanisms to regulate endogenous gene expression in cell nuclei has remained unclear.

### Changing miRISC complexes may affect gene expression

Regulating the composition and concentration of active miRISC complexes offers a mechanism that can globally modulate gene expression. The TNRC6 scaffolding proteins are essential to miRNA function because TNRC6, rather than Argonaute proteins or miRNAs, directly recruit the factors necessary for assembly of a large miRISC complex that carries out canonical mRNA decay (95). We have observed that TNRC6A protein expression is reduced after 5 days of growth with cell density near 200% or greater (**Fig. 3BC**), and the interaction between AGO2 and TNRC6A is also lost (**Fig. 3C**). AGO2 localization is similar in the cytoplasm and the nucleus after 5 days of growth with cell density close to 200% (**Fig. 1D**), yet we observe significant change in AGO2 localization to the nucleus only after 6-7 days of growth with cell density closer to 300-400% (**Fig. 1BCDE**). These results suggest that loss of TNRC6A expression may contribute to the shift of AGO2 localization to the nucleus (**Fig. 1D**, **Fig. 3B**).

We also observed that the constitutive deletion of TNRC6A and TNRC6B resulted in almost exclusive nuclear localization of AGO2 (**Fig. 3A**), and this suggests that TNRC6 proteins may function to anchor AGO in the cytoplasm. This supports previous studies that suggest TNRC6 expression is critical for determining the localization of AGO2 (45,47,69), composition of active high molecular weight RISC complexes and inactive low molecular weight RISC complexes (6,96-98), and miRNA activity (6,96-98).

### Implications of nuclear localization of AGO2 for cell growth, migration, and cancer

Compromised miRNA activity may provide specific advantages for cancer cells. miRNAs are globally depleted in tumors relative to their normal tissue counterparts (99). 3’UTR’s are frequently shortened in tumors via alternative polyadenylation site choice (100,101). The trafficking of AGO2 into the nucleus may be an additional mechanism for regulating miRNA activity.

If RNAi is a surveillance system to silence aberrant transcripts, what would happen if cells shuttle at least half of the Argonaute pool to the nucleus? Perhaps the remaining cytoplasmic Argonaute pool may be sufficient to sustain some miRISC regulation but insufficient to mediate the degradation of all aberrant transcripts, allowing for some oncogenic mRNAs to be translated and even upregulated.

Ebert and Sharp have noted the potential importance of miRNAs in buffering gene expression and increasing the robustness of normal physiologic processes (38). While individual miRNAs may have only modest effects on individual genes, the aggregate effect of all miRNAs on all genes might have a more significant impact. They pointed out that widespread de-repression of miRNA function could lead to unbuffering of gene expression (38). They note that widespread “unbuffering” of miRNA control might increase the heterogeneity of gene expression within a population of tumor cells. Increased heterogeneity might enhance the potential for tumor growth, chemoresistance, and metastasis.

Our data suggest that the sequestration of AGO2 to the nucleus is another mechanism for regulating gene expression. Decreased levels of AGO2 in the cytoplasm would tend to decrease levels of canonical miRNA-mediated repression of mRNA targets. Consistent with that hypothesis, we observe that, when AGO2 localizes to nuclei of over-confluent cells, genes that are likely candidates for regulation by miRNAs in the cytoplasm show increased expression (**Figure 6B**).

We observed that growing cells at high cell density causes upregulation of miRISC target genes enriched in pathways affecting cell signaling, transduction, and migration (**Figure 9A**). Such widespread changes are consistent with the hypothesis that shifting AGO protein to the nucleus “unbuffers” cytoplasmic gene expression and that these global changes affect significant biological processes. Such changes are related to cell signaling, cell-cell junction organization, and cell migration, and our scratch assay data showed that increasing the amount of AGO2 in cell nuclei had no effect on proliferation (**Figure 9B**) yet increased the rate of migration (**Figure 9CD**).

To further pursue the potential link to tumor formation and cancer cell migration, we used wild-type HCT116 cells to grow 3-D tumor spheroid cultures (**Figure 10A**). Tumor spheroid cultures are useful cancer models because they mimic some of the properties of solid tumors and provide a more accurate depiction of cancer biology than growth of cells in 2-D (86). We observed that AGO2 is almost entirely localized to the nucleus of HCT116 cells 3-D cultured as spheroids in a collagen matrix (**Figure 10B**). We tested the expression of top candidate miRISC target genes *IGF1R*, *OLFML2A*, and *NOTCH3* and observed similar increases (**Figure 10C-E**) in expression of these candidate target genes that are involved in migration pathways as had been observed in cells grown to high confluence (**Figure 4**) or cells expressing NLS-AGO2 (**Figure 7**).

The data from overconfluent 2-D cell culture and 3-D tumor spheroids encouraged us to obtain human tissue samples from colon cancer patients (**Figure 11A**). Most patients had observable AGO2 in cell nuclei and eight patient samples had an increase of nuclear relative to cytoplasmic AGO2 in malignant versus normal adjacent tissue (**Figure 11BC**). Our sample size is too small to draw definitive conclusions, but a shift in AGO2 towards nuclear localization was associated with Grade 2 or Grade 3 tumors.

We did not observe a simple or universal correlation between nuclear AGO2, cancer diagnosis, or cancer grade, even within the grade 2 and grade 3 samples. This may be due to the diverse natural histories of each sample. The variation in soluble nuclear and chromatin localization between patient samples may be related to differing genetic backgrounds, cancer grade, or prior treatments. The finding that AGO2 can shift into the nucleus of tumor cells in a subset of patients suggests that the regulation of miRNA activity by changing the localization of core RISC machinery may contribute to cellular processes within some tumors and should be considered in studies aimed at examining the impact of miRNA action in human cancers.

### Study Limitations

While we have evaluated effects of high cell density and nuclear localization of AGO2 on the control of genes likely to be regulated by miRNAs in the cytoplasm, we have not defined the potential of increased nuclear AGO to make a direct contribution to the control of transcription and splicing. We have examined only one cell line and its derivatives in depth, HCT116, although we do observe nuclear localization at high confluence in other cell lines (**Supplemental Fig 3**). It will be important to examine other types of cell stress and physiological conditions, extending the earlier studies of Scaziel and colleagues (41). The prevalence of AGO varies in different tissues (16,98) and the potential for nuclear RNAi in a cell or tissue type will need to be evaluated on a case-by-case basis. Finally, the small amount material available from individual human tumor samples limited the studies that could be performed for these initial investigations. We have not evaluated cellular AGO localization at the single cell level within the human tissues and it is possible that important effects may occur in specific sub-populations of cells.

### Conclusion

In this report, we use several model systems to show dynamic changes in relative localization of AGO2 between the cytoplasm and the nucleus. These include growing colorectal cancer cells to high confluence, depletion of miRNAs in the context of DROSHA knockout cells, rescue of miRNAs in the context of DROSHA knockout cells, depletion of TNRC6, expression of NLS-AGO2, expression of NES-AGO2, colorectal cancer cells grown in 3D culture to form tumor spheroids, and human colon tumor tissue (**Figure 12**). We find that there is not a static ratio of AGO2 between cell nuclei and cytoplasm. Multiple conditions allow AGO to “toggle” back and forth between nuclei and cytoplasm.

**Figure 12.**
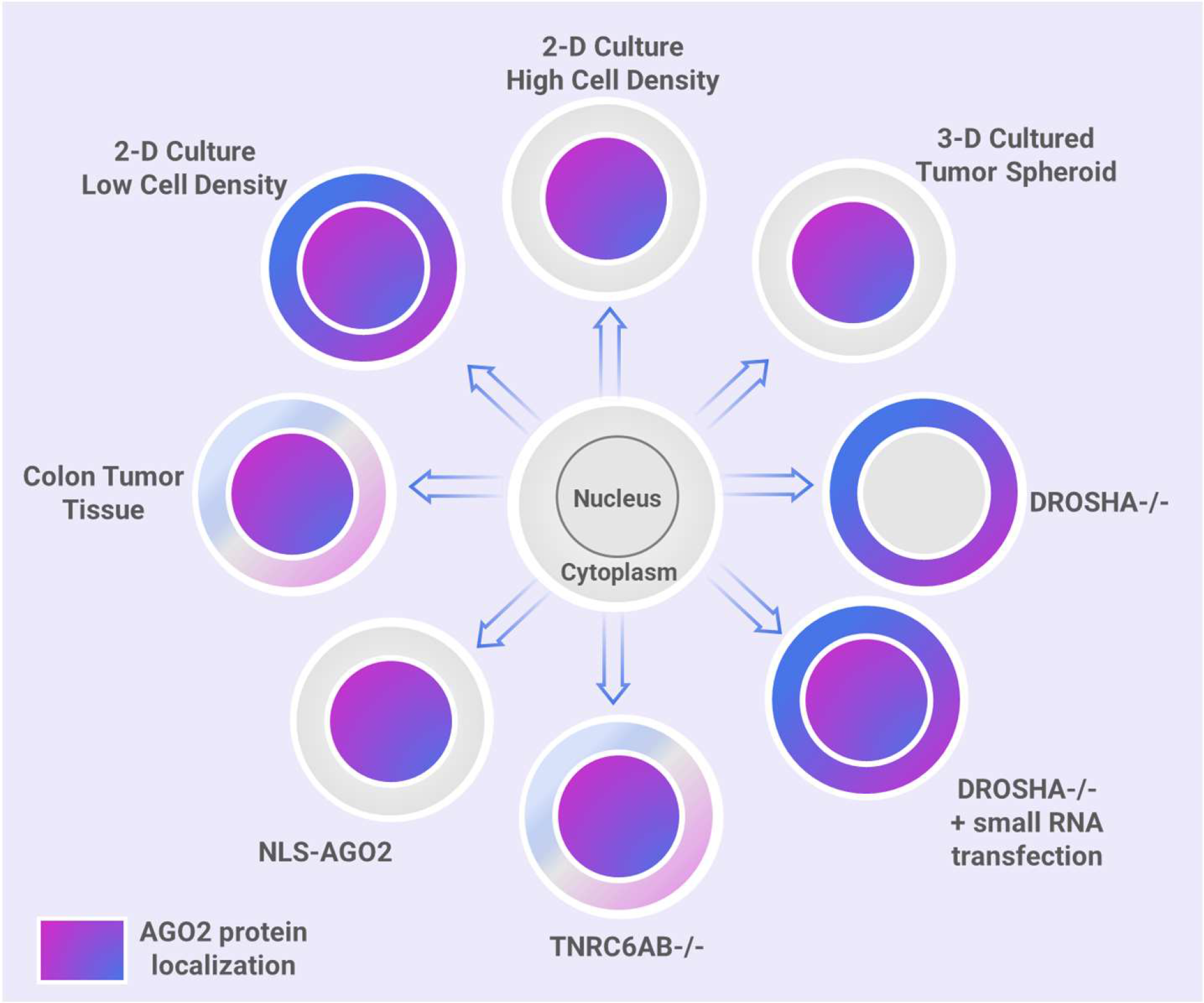
AGO2 subcellular localization depends on environmental conditions, availability of miRNAs, and expression of miRISC cofactor, TNRC6. Summary scheme showing that multiple variables can affect nuclear localization of AGO, including growth to high density in 2-D cell culture, knockout of DROSHA, knockout of DROSHA combined with introduction of exogenous small RNA, knockout of TNRC6, attachment of an NLS sequence to AGO, HCT116 3-D cultured to form tumor spheroids, and in human colon tumor tissue.

Our data are consistent with the conclusion that localizing AGO to cell nuclei de-represses genes that are likely to be controlled by canonical cytoplasmic RNAi. Our data are also consistent with a link between nuclear localization of AGO2 and cellular signaling related to positive regulation of cell migration. AGO2 nuclear localization is observed in tumor tissue from some patients, and the impact of shifting the localization of RNAi factors and reducing miRNA repression on cancer progression merits further exploration.

### Data Availability

The small-RNA and whole transcriptome RNA-sequencing data has been deposited in NCBI’s Gene Expression Omnibus (102) and is accessible through GEO Series accession number GSE236946. AGO2-eCLIP sequencing data was previously published (33).

## ACKNOWLEDGEMENTS

The authors thank the UTSW Genomics Sequencing Core for microRNA-seq library preparation, sequencing, and data analysis. We also thank the services of the Simmons Cancer Center’s Tissue Management Shared Resource, and the research reported in this publication was supported by the National Cancer Institute of the National Institutes of Health under award number 5P30CA142543. Microscopy reported in this publication was supported by the National Institute of Health under award number NIH 1S10OD028630-01 to Dr. Kate Luby-Phelps which was used to purchase the Nikon CSU-W1 with SoRa (Spinning disk confocal microscope). We thank Dr. Nicholas Conrad for critical advice on this manuscript.

## Conflict of interest statement

DRC is an Executive Editor of *Nucleic Acids Research*.

## FUNDING

This study was supported by 1F31GM137591 (K.C.J.), R35GM118103 (D.R.C.), and RO1CA243577 (R.A.B.) from the National Institutes of Health and the Robert A. Welch Foundation I-1244 (D.R.C.).

## Notes

### Competing Interest Statement

The authors have declared no competing interest.

### Summary of Updates

This version corrects the figure legend text.

