## Supplemental Data for "Nuclear Localization of Argonaute is affected by Cell Density and May Relieve Repression by microRNAs"

### Supplemental Figures

| Gene | Target | Primer Sequence 5’-3’ |
|---|---|---|
| IGF1R | mRNA | F: TTCTGGACAAGCCAGACAAC<br>R: CTCCTCTTTGATGCTGCTGAT |
| IGF1R | Pre-mRNA | F: AGACAACTGTCCTGACATGC<br>R: TCCTAATCTCCTGTGACCCAA |
| OLFML2A | mRNA | F: CGACAGGATCTATGTCACCAAC<br>R: CTGTGCCGATCCAGTTGTAG |
| NOTCH3 | mRNA | F: GCCAATAAGGACATGCAGGATAG<br>R: TGATCTCACGGTTGGCAAAG |
| RREB1 | mRNA | F: GGCAGCTTGATAACACAAAGAAA<br>R: CCCTGCTGTAGCTATCATTCAC |
| RREB1 | Pre-mRNA | F: AAGGAGGGCAGCTTGATAAC<br>R: TGACAATTAAACGGGAGGTAGG |
| RNF24 | mRNA | F: CTTCAAGCCTCGAGATGAGTT<br>R: GGGACACACTTTACGAACCT |
| RNF24 | Pre-mRNA | F: GAAGACTTCAAGCCTCGAGATG<br>R: CTCAAACTCGGGTCCCATTAAG |
| VEGFA | mRNA | F: CGAGTACATCTTCAAGCCATCC<br>R: GGTGAGGTTTGATCCGCATAAT |
| VEGFA | Pre-mRNA | F: GAGGGCAGAATCATCACGAA<br>R: AATTAGGCCATCCACCCATC |
| HPRT1 | mRNA | F: AGTTCTGTGGCCATCTGCTTAGTAG<br>R: AAACAACAATCCGCCCAAAGG |
| snRNP200 | mRNA | F: CGAGAAGTGGGACATCATCAC<br>R: ACCTCTGTCATCGTGGAGAA |
| ZMYM4 | mRNA | F: GAGGGAAAGTGGAAGAGTTCTG<br>R: CTGAGTTTACCCTGTCGCTTAC |
| COL6A1 | mRNA | F: CGCTGGTCAAGGAGAACTATG<br>R: CAGGTGTAATCTGGACACTTCTT |
| PRDM1 | mRNA | F: GTATGCCACCAACAGTGAAGA<br>R: AGGAACTGTGTCATTGGTGTAG |
| MINK1 | mRNA | F: GGCTCAAGTTCCTGTGTGAG<br>R: TCACCAGTTCATGATGCAGTTA |
| SERPINB9 | mRNA | F: CTGGTGCTGCTGCCTGA<br>R: GGAGAACTTCAACCTCAGTACTCTTC |

**Supplemental Table 1. Sequences of primers used to test expression of candidate miRISC target genes and internal control genes.**

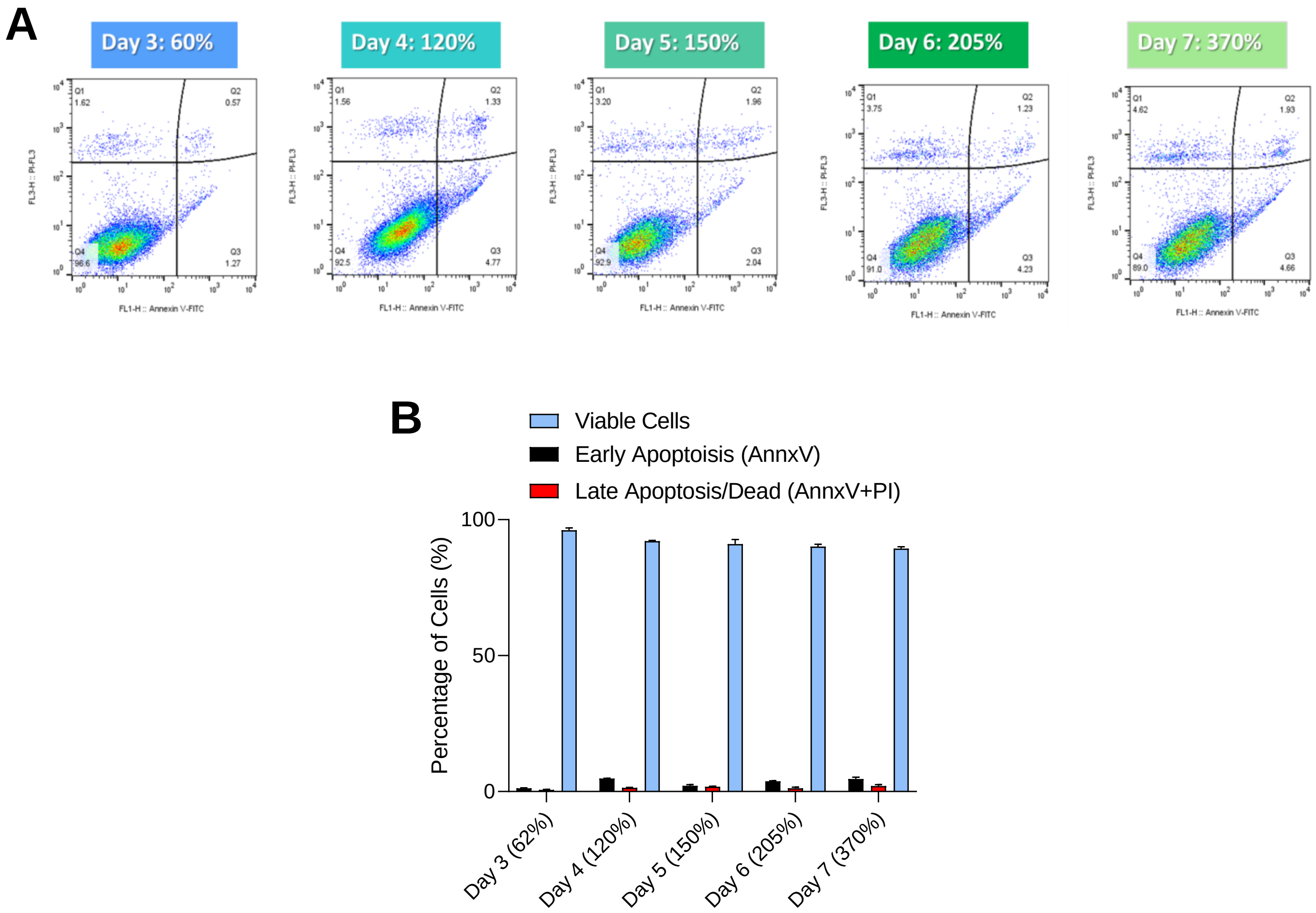


**Supplemental Figure 1. Majority of HCT116 cells are viable throughout high cell density growth conditions.**

(A) Scatter plots of flow cytometry data from cells grown to high cell density. Three biological replicates were performed for each cell density time point, and one representative plot is shown per day. (B) Bar graph summarizing dual-viability staining results in (A) with percentage of viable cells in blue, early apoptotic cells in black, and late apoptotic/dead cells in red. Statistical significance calculated using multiple paired t-tests relative to Day 3. HCT116 cells stained with Annexin V-FITC (AnnxV) and propidium iodide (PI) after growing for a total of seven days with Day 3-7 shown.

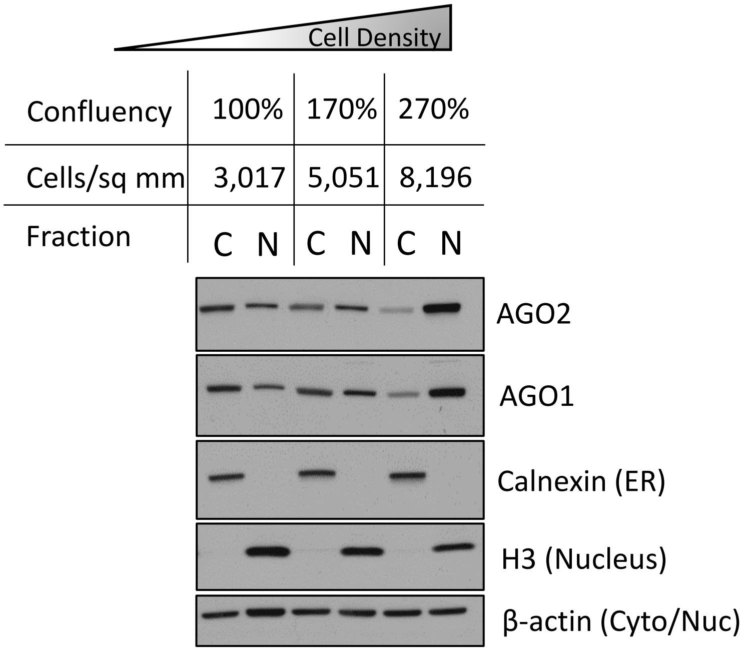


**Supplemental Figure 2. AGO1 and AGO2 localization as a function of cell density.** Representative Western blot showing AGO1 and AGO2 localization as a function of cell density. Same cell lysate samples as in Figure 1D with AGO1 localization shown.

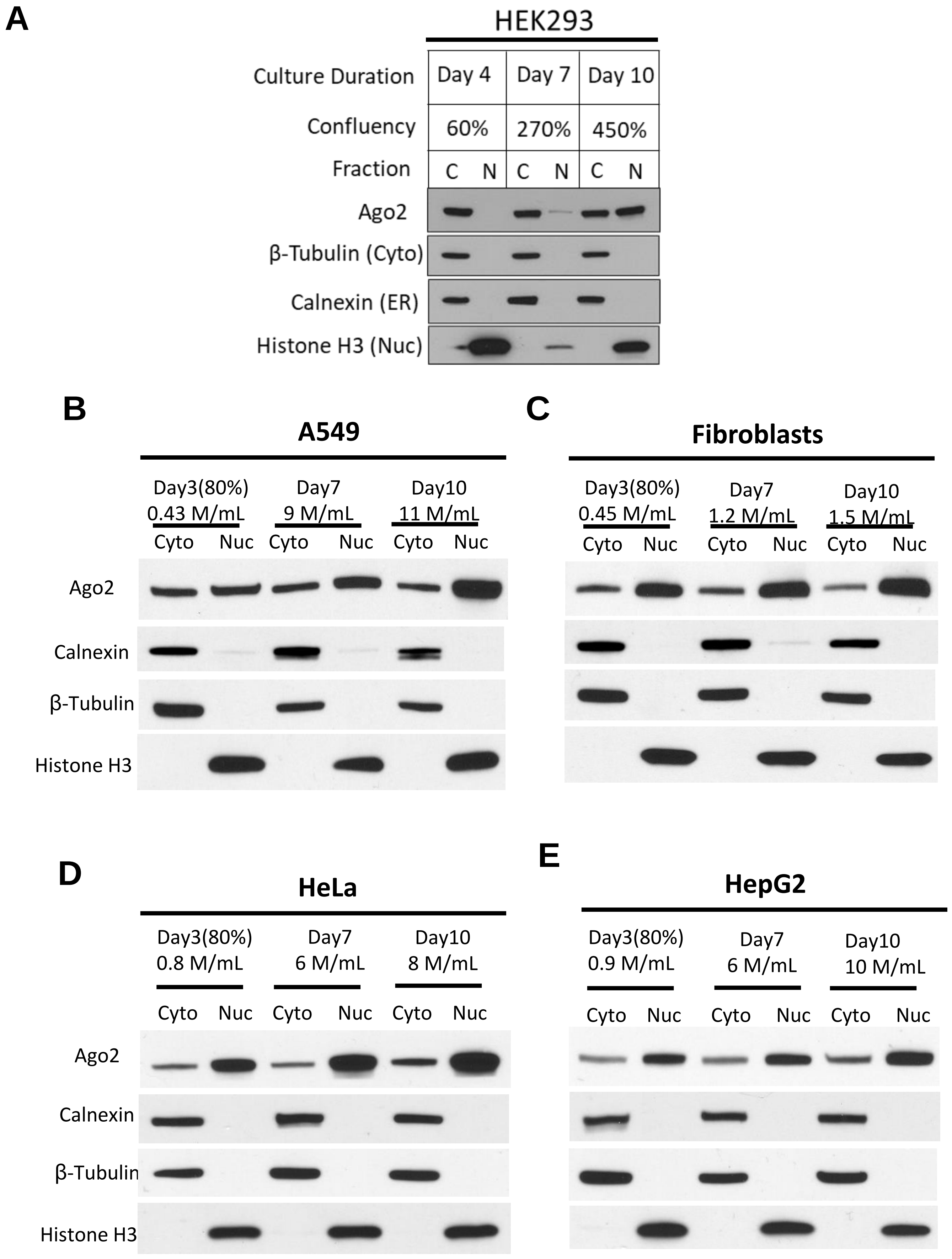


**Supplemental Figure 3. AGO2 localization in other cell lines.** Western blot showing AGO2 localization of different cell lines grown to high cell density. (A) HEK293, (B) A549, (C) fibroblasts, which do not grow beyond a monolayer, (D) HeLa, and (E) HepG2.

**A**

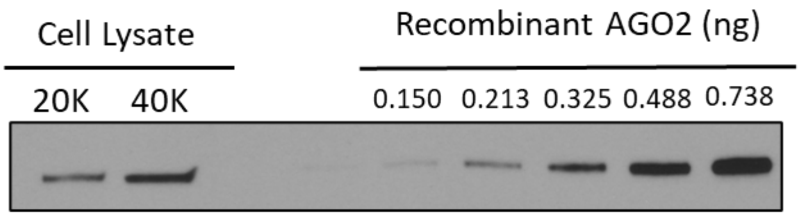


| Lane | Signal | Mass Calculation | Cells | AGO2 Molecules | AGO2 Copy per cell |
|---|---|---|---|---|---|
| 1 | 12060.539 | 0.274464815 | 20,000 | 1.65E+09 | 82,614 |
| 2 | 24734.731 | 0.471869837 | 40,000 | 2.84E+09 | 71,016 |
| | | | | **Average** | **76,815** |

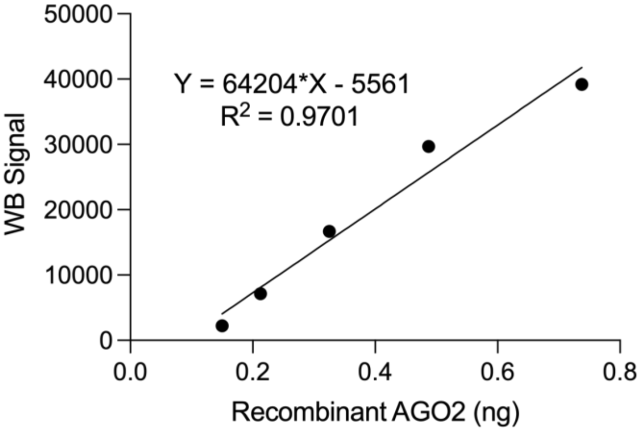


**B**

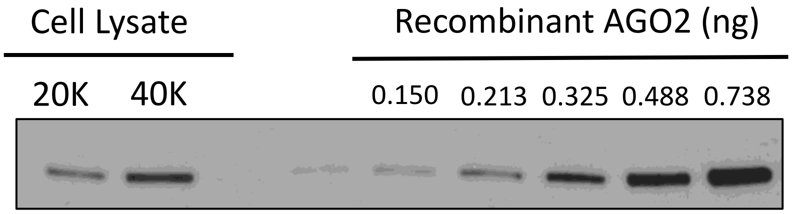


| Lane | Signal | Mass Calculation | Cells | AGO2 Molecules | AGO2 Copy per cell |
|---|---|---|---|---|---|
| 1 | 8683.468 | 0.199837 | 20,000 | 1.20E+09 | 60,151 |
| 2 | 21690.075 | 0.392359 | 40,000 | 2.36E+09 | 59,050 |
| | | | | **Average** | **59,600** |

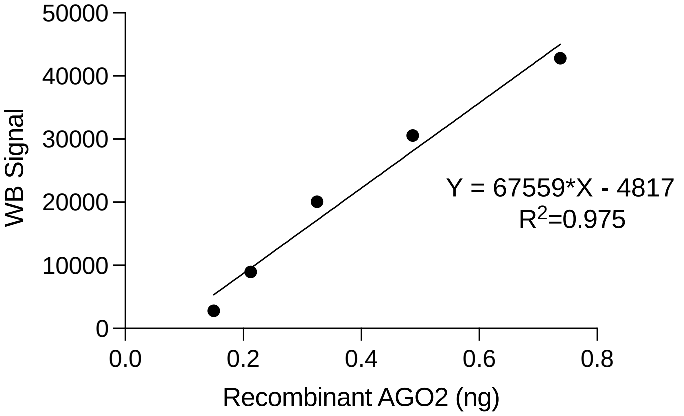


**C**

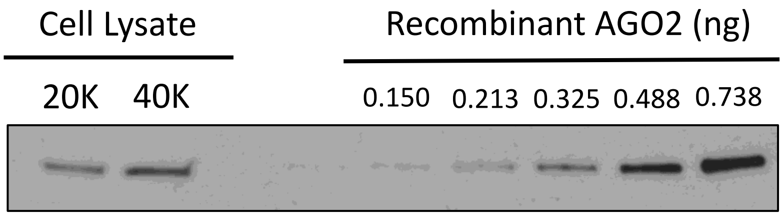


| Lane | Signal | Mass Calculation | Cells | AGO2 Molecules | AGO2 Copy per cell |
|---|---|---|---|---|---|
| 1 | 12203.711 | 0.298088 | 20,000 | 1.79E+09 | 89,724 |
| 2 | 22266.368 | 0.427861 | 40,000 | 2.58E+09 | 64,393 |
| | | | | **Average** | **77,059** |

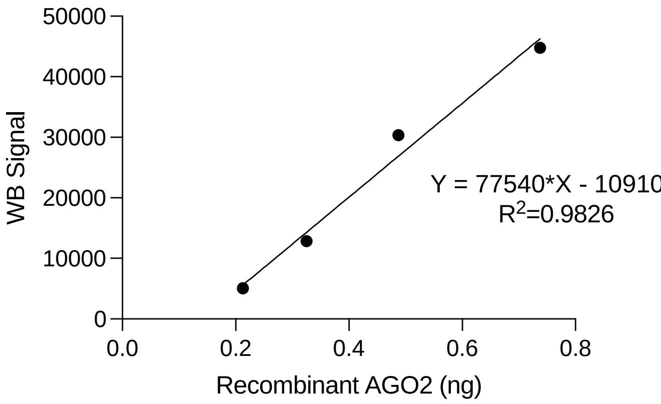


**Supplementary Figure 4.** (A-C) Western blots, linear regressions and calculated mass and copy number of AGO2 for <u>Day 3 whole cell</u> absolute quantification. Densitometry was used to produce linear regressions from a serial dilution of recombinant AGO2. Mass of AGO2 per lysate sample was calculated based on linear regression and AGO2 molecular weight and number of cells loaded per lysate were used to calculate number of AGO2 molecules and copy number per cell.

**D**

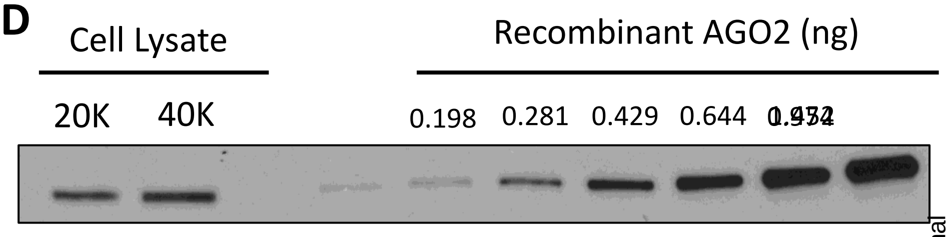


| Lane | Signal | Mass Calculation | Cells | AGO2 Molecules | AGO2 Copy per cell |
|---|---|---|---|---|---|
| 1 | 14351.832 | 0.26888 | 20,000 | 1.62E+09 | 80,933 |
| 2 | 21457.075 | 0.475403 | 40,000 | 2.86E+09 | 71,548 |
| | | | | **Average** | **76,240** |

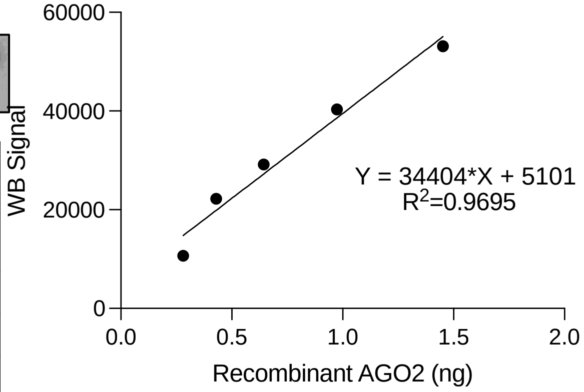


**E**

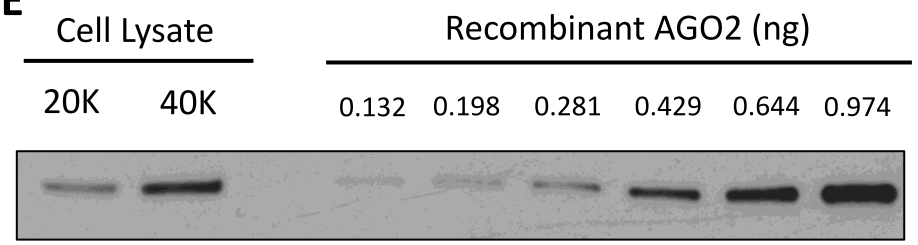


| Lane | Signal | Mass Calculation | Cells | AGO2 Molecules | AGO2 Copy per cell |
|---|---|---|---|---|---|
| 1 | 8457.175 | 0.257588 | 20,000 | 1.55E+09 | 77,534 |
| 2 | 25794.61 | 0.595846 | 40,000 | 3.59E+09 | 89,675 |
| | | | | **Average** | **83,604** |

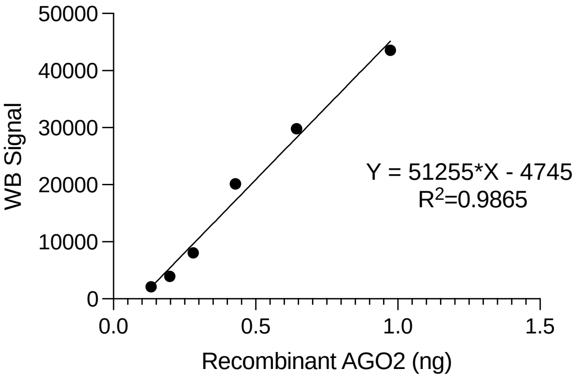


**F**

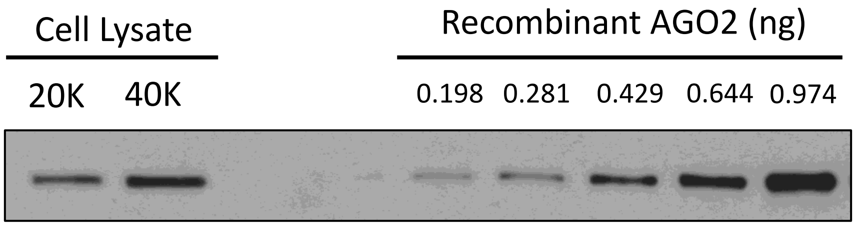


| Lane | Signal | Mass Calculation | Cells | AGO2 Molecules | AGO2 Copy per cell |
|---|---|---|---|---|---|
| 1 | 13455.711 | 0.373897 | 20,000 | 2.25E+09 | 112,543 |
| 2 | 26633.024 | 0.641039 | 40,000 | 3.86E+09 | 96,476 |
| | | | | **Average** | **104,510** |

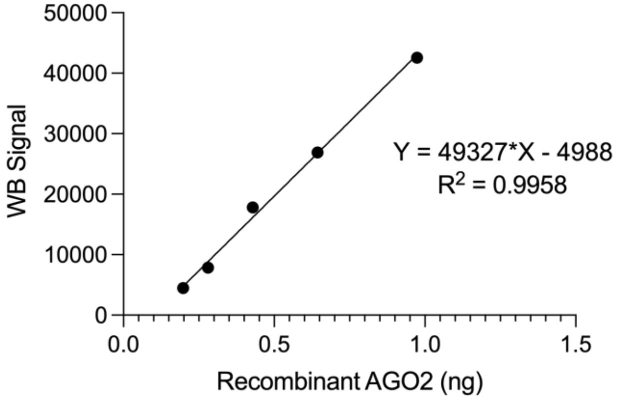


**Supplementary Figure 4.** (D-F) Western blots, linear regressions and calculated mass and copy number of AGO2 for Day 7 whole cell absolute quantification. Densitometry was used to produce linear regressions from a serial dilution of recombinant AGO2. Mass of AGO2 per lysate sample was calculated based on linear regression and AGO2 molecular weight and number of cells loaded per lysate were used to calculate number of AGO2 molecules and copy number per cell.

**G**

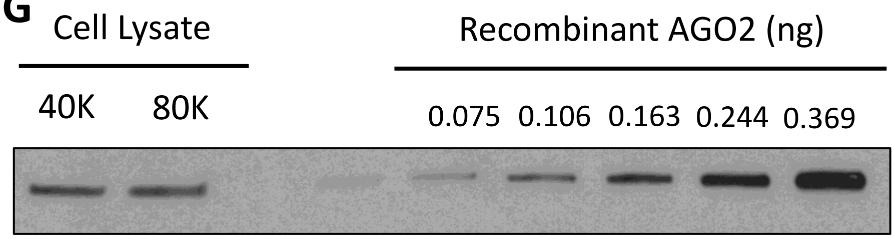


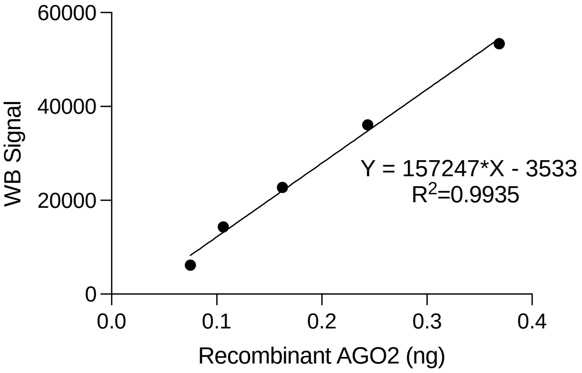


| Lane | Signal | Mass Calculation | Cells | AGO2 Molecules | AGO2 Copy per cell |
|---|---|---|---|---|---|
| 1 | 23387.874 | 0.171201 | 40,000 | 1.03E+09 | 25,766 |
| 2 | 24344.267 | 0.177283 | 80,000 | 1.07E+09 | 13,341 |
| | | | | **Average** | **19,553** |

**H**

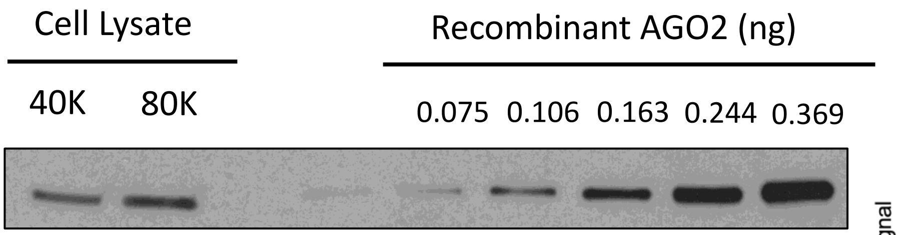


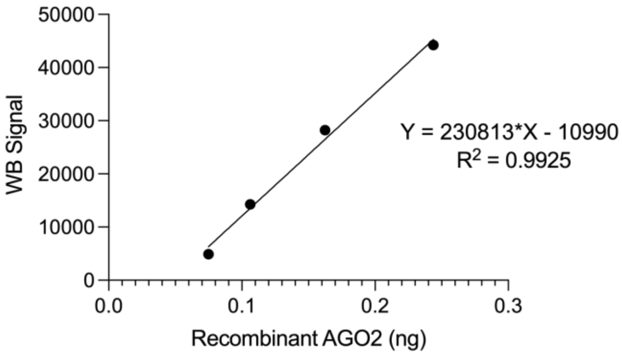


| Lane | Signal | Mass Calculation | Cells | AGO2 Molecules | AGO2 Copy per cell |
|---|---|---|---|---|---|
| 1 | 16558.61 | 0.119355 | 40,000 | 7.19E+08 | 17,963 |
| 2 | 26173.388 | 0.161011 | 80,000 | 9.69E+08 | 12,116 |
| | | | | **Average** | **15,039** |

**I**

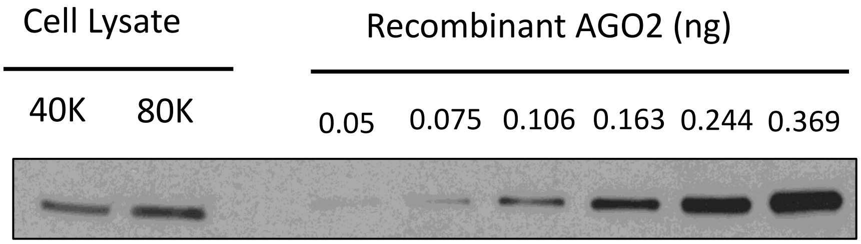


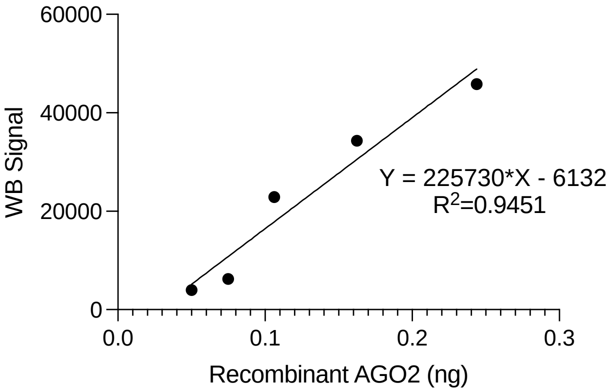


| Lane | Signal | Mass Calculation | Cells | AGO2 Molecules | AGO2 Copy per cell |
|---|---|---|---|---|---|
| 1 | 14299.075 | 0.091322 | 40,000 | 5.50E+08 | 13,744 |
| 2 | 16598.267 | 0.101599 | 80,000 | 6.12E+08 | 7,645 |
| | | | | **Average** | **10,695** |

**Supplementary Figure 4.** (G-I) Western blots, linear regressions and calculated mass and copy number of AGO2 for Day 3 cytoplasmic fraction absolute quantification. Densitometry was used to produce linear regressions from a serial dilution of recombinant AGO2. Mass of AGO2 per lysate sample was calculated based on linear regression and AGO2 molecular weight and number of cells loaded per lysate were used to calculate number of AGO2 molecules and copy number per cell.

**J**

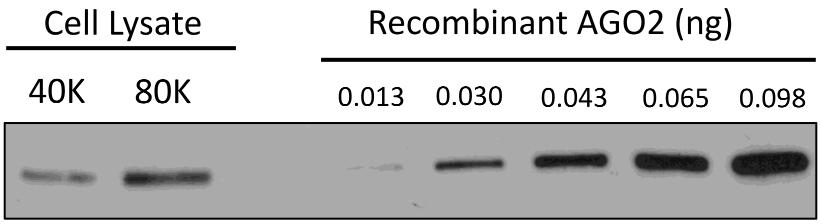


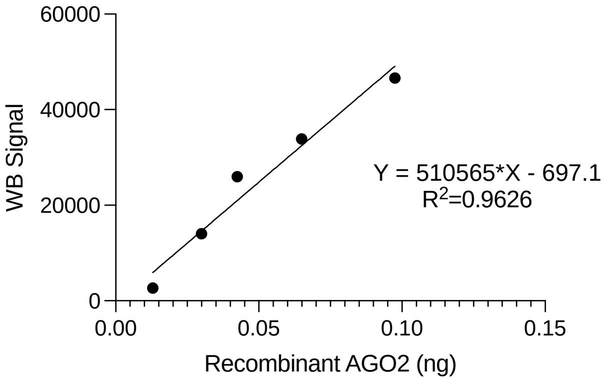


| Lane | Signal | Mass Calculation | Cells | AGO2 Molecules | AGO2 Copy per cell |
|---|---|---|---|---|---|
| 1 | 10477.024 | 0.021886 | 40,000 | 1.32E+08 | 3,294 |
| 2 | 24615.288 | 0.049577 | 80,000 | 2.98E+08 | 3,731 |
| | | | | **Average** | **3,512** |

**K**

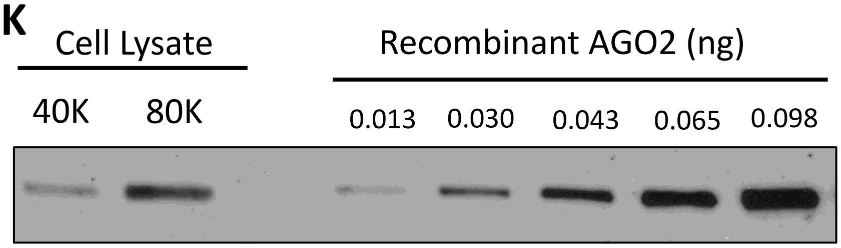


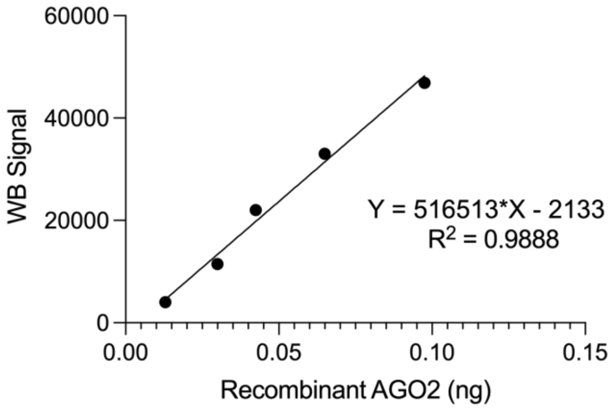


| Lane | Signal | Mass Calculation | Cells | AGO2 Molecules | AGO2 Copy per cell |
|---|---|---|---|---|---|
| 1 | 7529.903 | 0.018709 | 40,000 | 1.13E+08 | 2,816 |
| 2 | 24938.095 | 0.052412 | 80,000 | 3.16E+08 | 3,944 |
| | | | | **Average** | **3,380** |

**L**

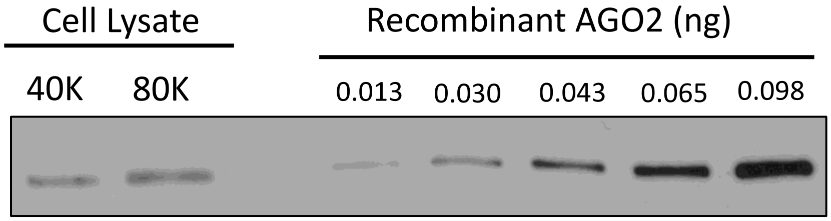


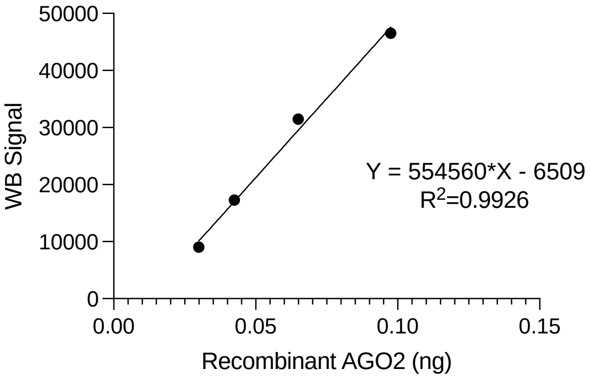


| Lane | Signal | Mass Calculation | Cells | AGO2 Molecules | AGO2 Copy per cell |
|---|---|---|---|---|---|
| 1 | 8439.418 | 0.026955 | 40,000 | 1.62E+08 | 4,057 |
| 2 | 15942.631 | 0.040485 | 80,000 | 2.44E+08 | 3,046 |
| | | | | Average | 3,552 |

**Supplementary Figure 4.** (J-L) Western blots, linear regressions and calculated mass and copy number of AGO2 for Day 7 cytoplasmic fraction absolute quantification. Densitometry was used to produce linear regressions from a serial dilution of recombinant AGO2. Mass of AGO2 per lysate sample was calculated based on linear regression and AGO2 molecular weight and number of cells loaded per lysate were used to calculate number of AGO2 molecules and copy number per cell.

**M**

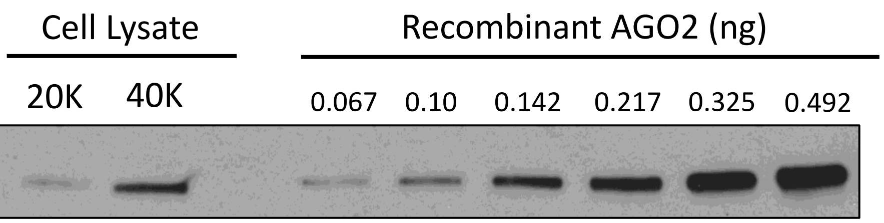


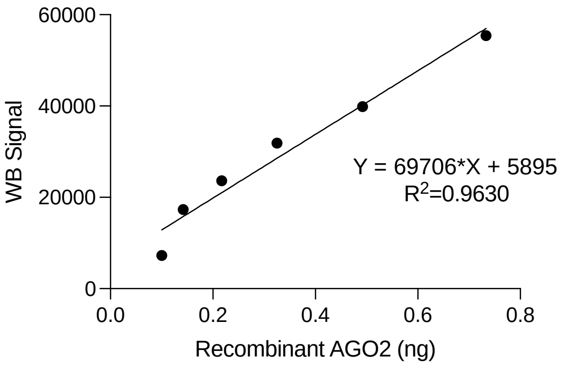


| Lane | Signal | Mass Calculation | Cells | AGO2 Molecules | AGO2 Copy per cell |
|---|---|---|---|---|---|
| 2 | 16213.146 | 0.171872 | 40,000 | 1.03E+09 | 25,867 |
| | | | | **Average** | **25,867** |

**N**

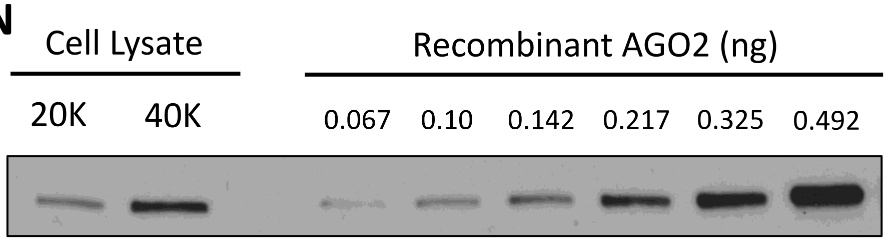


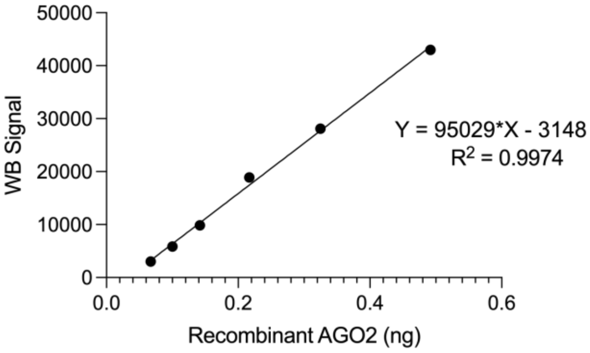


| Lane | Signal | Mass Calculation | Cells | AGO2 Molecules | AGO2 Copy per cell |
|---|---|---|---|---|---|
| 1 | 6572.882 | 0.102296 | 20,000 | 6.16E+08 | 30,791 |
| 2 | 21230.075 | 0.256535 | 40,000 | 1.54E+09 | 38,609 |
| | | | | **Average** | **34,700** |

**O**

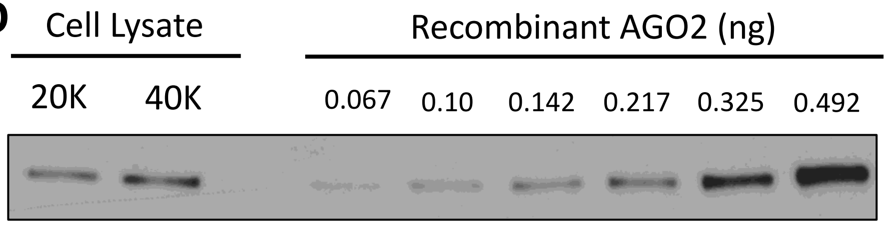


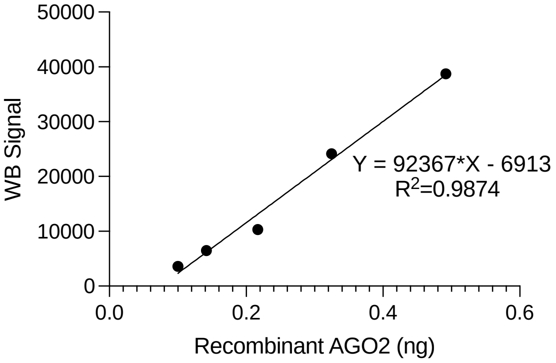


| Lane | Signal | Mass Calculation | Cells | AGO2 Molecules | AGO2 Copy per cell |
|---|---|---|---|---|---|
| 1 | 7910.711 | 0.16049 | 20,000 | 9.66E+08 | 48,308 |
| 2 | 15390.539 | 0.24147 | 40,000 | 1.45E+09 | 36,341 |
| | | | | **Average** | **42,324** |

**Supplementary Figure 4.** (M-O) Western blots, linear regressions and calculated mass and copy number of AGO2 for day 3 nuclear fraction absolute quantification. Densitometry was used to produce linear regressions from a serial dilution of recombinant AGO2. Mass of AGO2 per lysate sample was calculated based on linear regression and AGO2 molecular weight and number of cells loaded per lysate were used to calculate number of AGO2 molecules and copy number per cell.

**P**

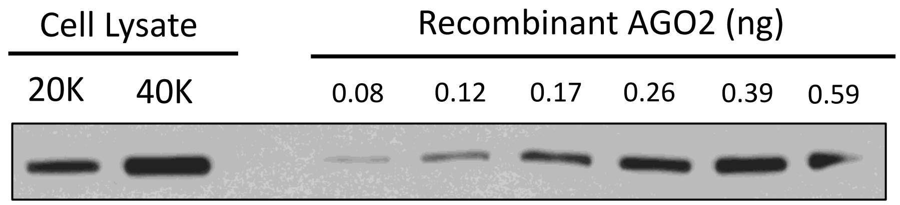


| Lane | Signal | Mass Calculation | Cells | AGO2 Molecules | AGO2 Copy per cell |
|---|---|---|---|---|---|
| 1 | 40199.338 | 0.231832 | 20,000 | 1.40E+09 | 69,781 |
| 2 | 74011.016 | 0.386894 | 40,000 | 2.33E+09 | 58,228 |
| | | | | **Average** | **64,004** |

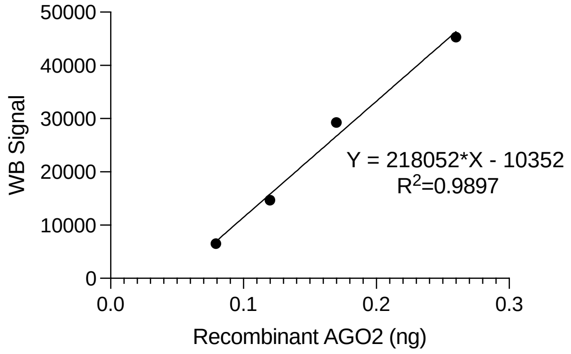


**Q**

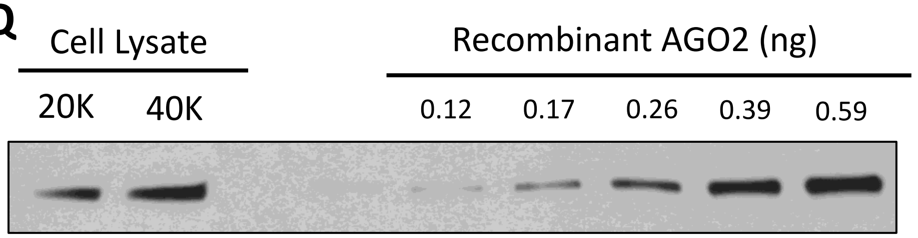


| Lane | Signal | Mass Calculation | Cells | AGO2 Molecules | AGO2 Copy per cell |
|---|---|---|---|---|---|
| 1 | 18464.418 | 0.260777 | 20,000 | 1.57E+09 | 78,494 |
| 2 | 34665.439 | 0.436065 | 40,000 | 2.63E+09 | 65,628 |
| | | | | **Average** | **72,061** |

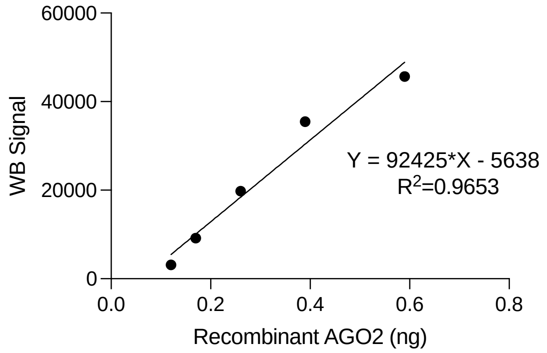


**R**

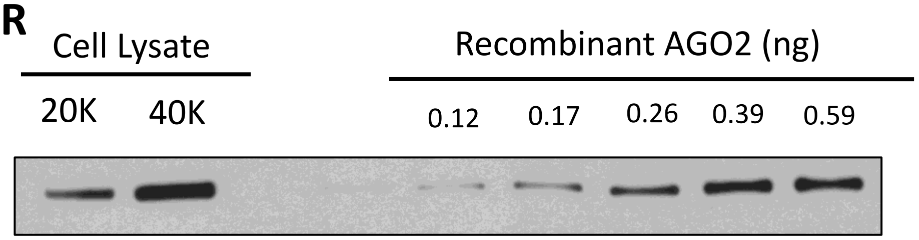


| Lane | Signal | Mass Calculation | Cells | AGO2 Molecules | AGO2 Copy per cell |
|---|---|---|---|---|---|
| 1 | 26987.731 | 0.277917 | 20,000 | 1.67E+09 | 83,653 |
| 2 | 63852.167 | 0.550536 | 40,000 | 3.31E+09 | 82,856 |
| | | | | **Average** | **83,254** |

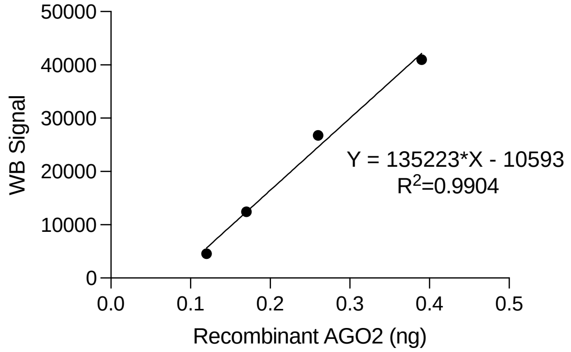


**Supplementary Figure X.** (P-R) Western blots, linear regressions and calculated mass and copy number of AGO2 for day 7 nuclear fraction absolute quantification. Densitometry was used to produce linear regressions from a serial dilution of recombinant AGO2. Mass of AGO2 per lysate sample was calculated based on linear regression and AGO2 molecular weight and number of cells loaded per lysate were used to calculate number of AGO2 molecules and copy number per cell.

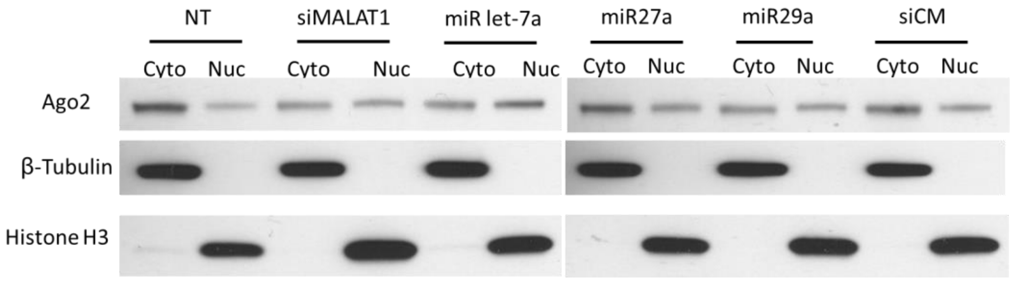


**Supplementary Figure 5. AGO2 nuclear localization is sensitive to microRNA depletion**. (A) Western blot showing transfection of microRNA mimics and control oligos into Drosha-/- cells rescues nuclear AGO2 localization.

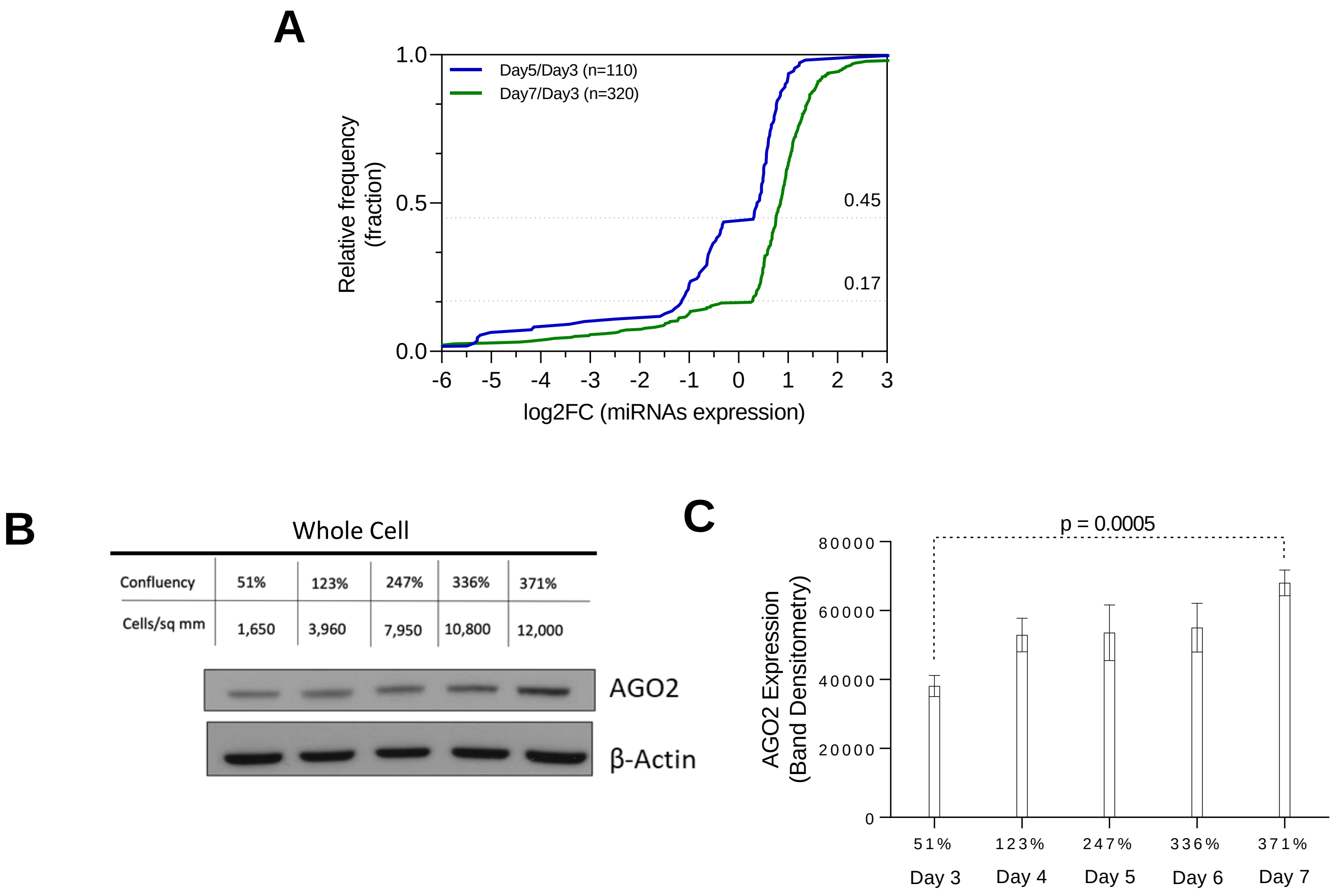


**Supplemental Figure 6. Global miRNA expression increase matches increase in total AGO2 protein levels.** (A) Cumulative distribution frequency plot showing that over 80% of significantly effected miRNAs by Day 7 are upregulated. (B) Western blot showing AGO2 expression in whole cell HCT116 at increasing cell density. (C) Quantification of AGO2 band densitometry relative to β-Actin in (B). Increase in AGO2 is significant on Day 7 relative to Day 3. Unpaired t-test with Welch's correction to calculate p-value of 0.0005.

**A**

| miRNA Rank by Abundance | Total miRNA Family Seeds: 8 | Cyto-3'UTR-No6mer Total AGO2 Binding Sites: 29 | Seed | log2FC(Day7/Day3) | padj |
|---|---|---|---|---|---|
| #1 | miR-10 | IGF1R | 7mer-a1 | 2.07 | 2.9E-124 |
| | | IGF1R | 7mer-m8 | | |
| | | IGF1R | 7mer-a1 | | |
| | | IGF1R | 8mer | | |
| | | IGF1R | 7mer-a1 | | |
| | | IGF1R | 8mer | | |
| | | IGF1R | 8mer | | |
| #3 | let-7a | IGF1R | 7mer-a1 | | |
| | | IGF1R | 7mer-a1 | | |
| | | IGF1R | 7mer-a1 | | |
| | | IGF1R | 7mer-a1 | | |
| | | IGF1R | 7mer-a1 | | |
| | | IGF1R | 7mer-a1 | | |
| | | IGF1R | 7mer-a1 | | |
| | | IGF1R | 7mer-a1 | | |
| | | IGF1R | 7mer-a1 | | |
| #8 | miR-22 | IGF1R | 7mer-a1 | | |
| #9 | miR-27 | IGF1R | 7mer-a1 | | |
| #10 | miR-182 | IGF1R | 7mer-a1 | | |
| #13 | miR-15 | IGF1R | 7mer-m8 | | |
| | | IGF1R | 7mer-m8 | | |
| | | IGF1R | 7mer-m8 | | |
| | | IGF1R | 7mer-m8 | | |
| | | IGF1R | 7mer-m8 | | |
| | | IGF1R | 7mer-m8 | | |
| | | IGF1R | 7mer-m8 | | |
| | | IGF1R | 7mer-m8 | | |
| #16 | miR-17 | IGF1R | 7mer-a1 | | |
| #17 | miR-181 | IGF1R | 8mer | | |

**B**

| miRNA Rank by Abundance | Total miRNA Family Seeds: 1 | Cyto-3'UTR-No6mer Total AGO2 Binding Sites: 2 | Seed | log2FC(Day7/Day3) | padj |
|---|---|---|---|---|---|
| #9 | miR-27 | NOTCH3 | 7mer-a1 | 3.88 | 3.5E-73 |
| | | NOTCH3 | 7mer-m8 | | |

**C**

| miRNA Rank by Abundance | Total miRNA Family Seeds: 2 | Cyto-3'UTR-No6mer Total AGO2 Binding Sites: 2 | Seed | log2FC(Day7/Day3) | padj |
|---|---|---|---|---|---|
| #10 | miR-182 | OLFML2A | 7mer-a1 | 2.60 | 4.6E-81 |
| #12 | miR-29 | OLFML2A | 7mer-m8 | | |

**Supplemental Figure 7. miRNAs with seed matches within regions of AGO2 binding sites for top three candidate miRISC targets.** (A-C) microRNAs that have binding sites within regions of AGO2 occupancy from AGO2-eCLIP-sequencing within (A) IGF1R, (B) NOTCH3, and (C) OLFML2A 3’UTR in cytoplasm.

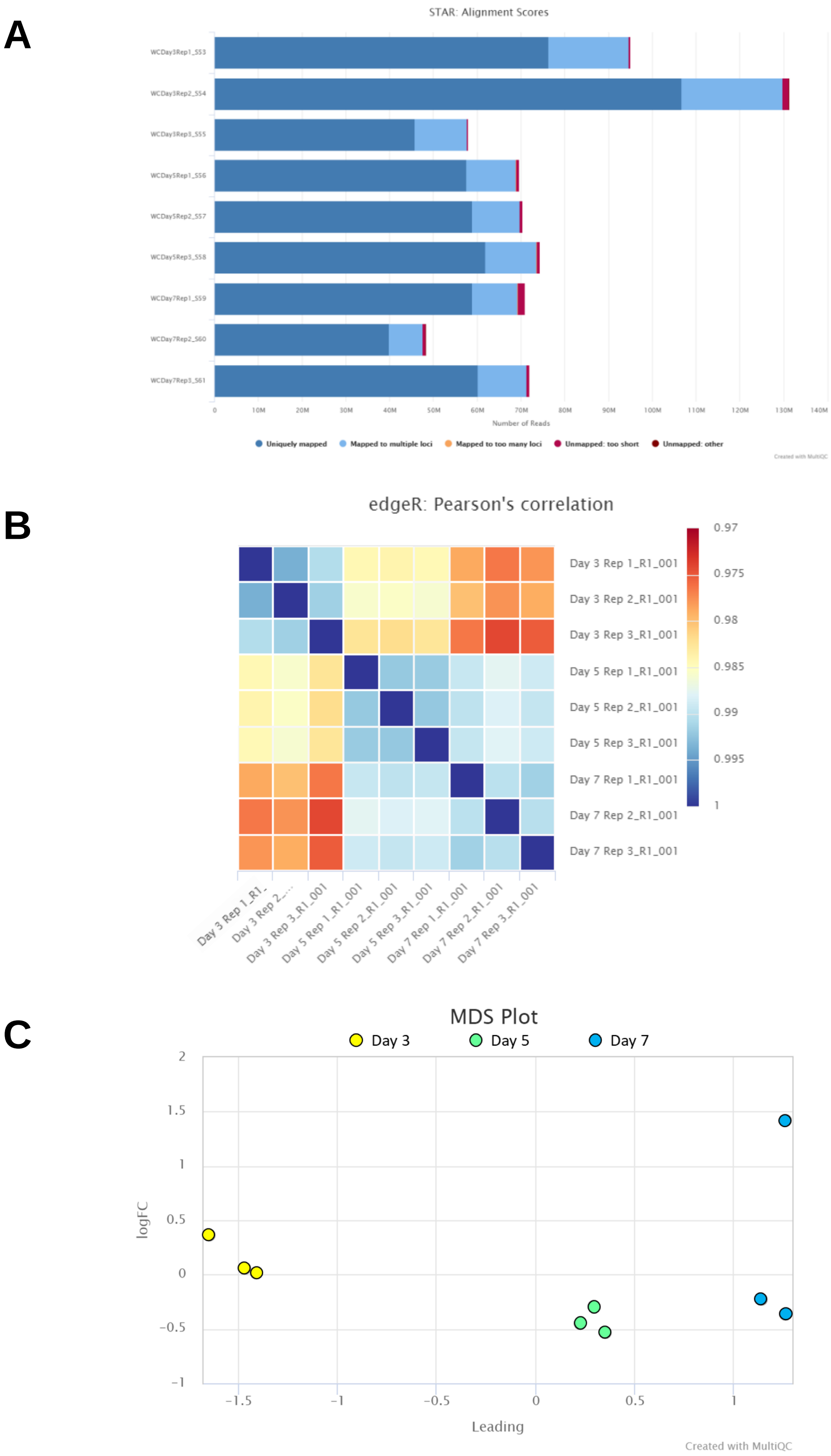


**Supplemental Figure 8. Whole Transcriptome Sequencing Quality Control.** (A) STAR Alignment Scores describing unique reads. (B) edgeR: Sample Similarity is generated from normalized gene counts through edgeR. Pearson's correlation between log2 normalized CPM values are then calculated and clustered. (C) Multidimensional scaling (MDS) plot showing clustering of each replicate submitted for whole transcriptome sequencing. (A-C) Plots generated by MultiQC.

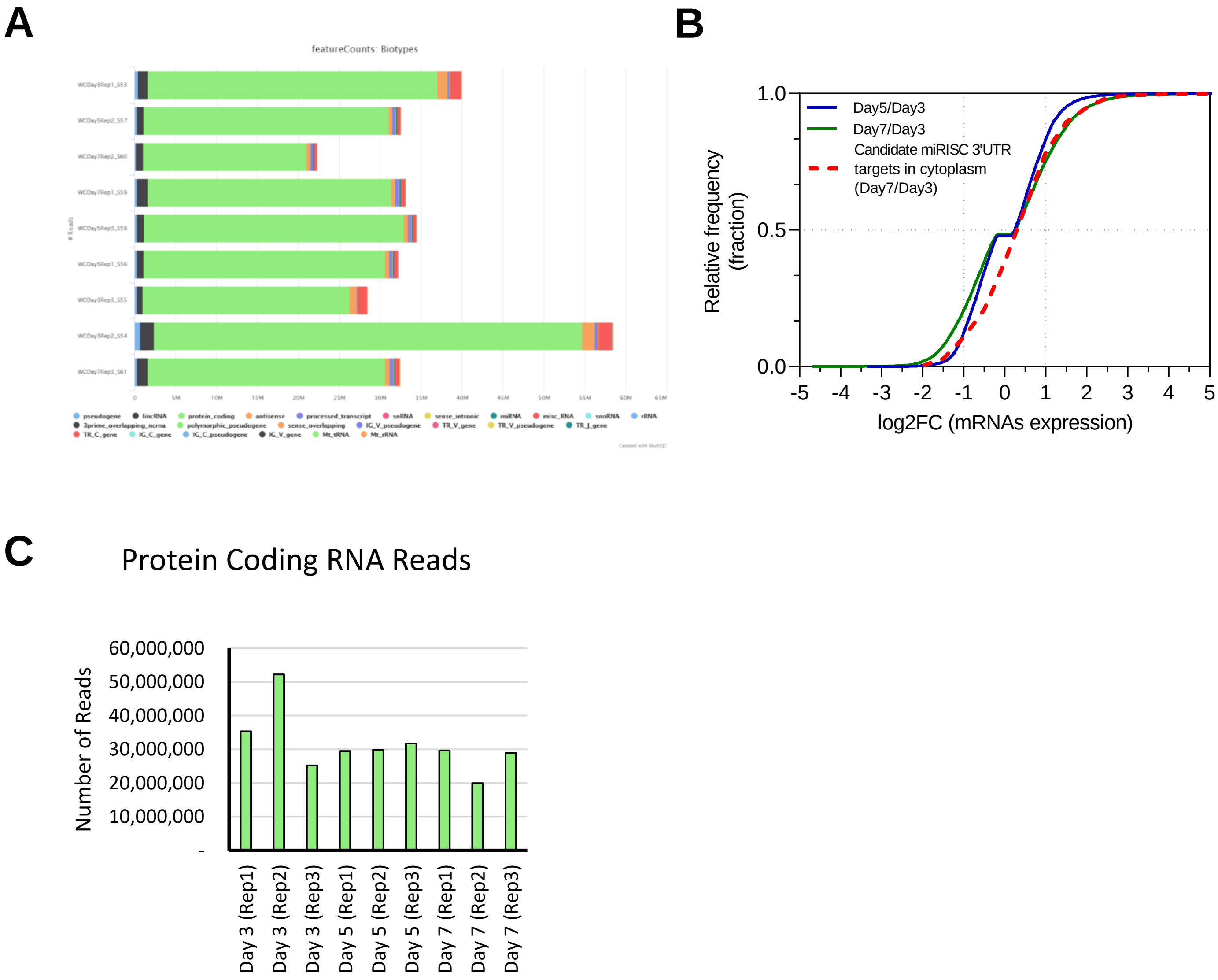


**Supplemental Figure 9.** (A) Biotype Counts, from whole transcriptome RNA-sequencing analysis, show reads overlapping genomic features of different biotypes, counted by featureCounts**.** (B) Cumulative distribution frequency plot showing global differential expression on Day 5 relative to Day 3 (blue), Day 7 relative to Day 3 (green), and the subset of candidate cytoplasm miRISC targets (dotted red line, all genes from Fig 5B). (C) Biotype counts from (A) showing reads aligning to protein coding RNAs.

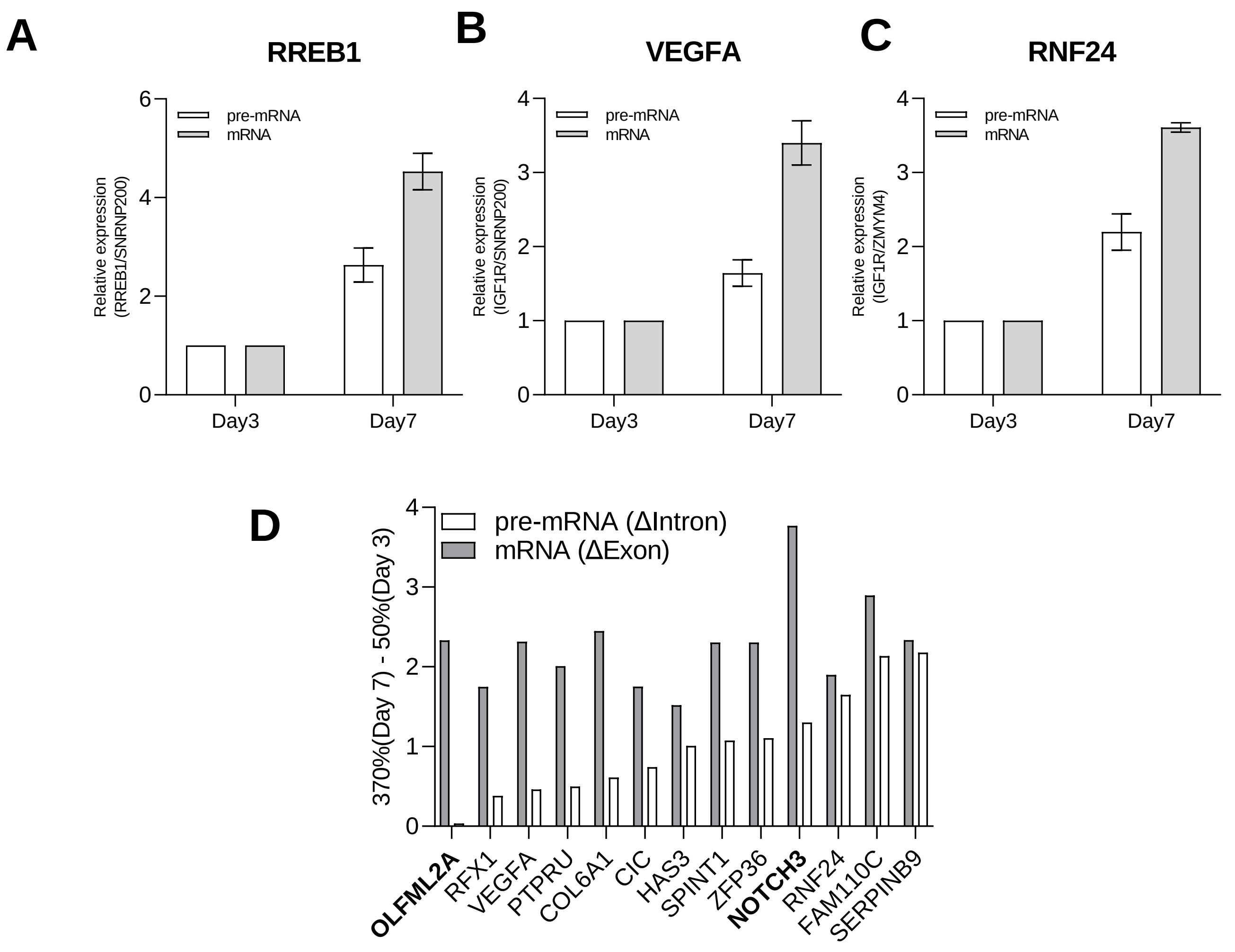


**Supplemental Figure 10.** (A-C) RT-qPCR measurements of relative expression of pre-mRNA and mRNA for candidate cytoplasmic RISC targets on Day 7 (370% confluent) relative to Day 3 (50% confluent). (D) Exon Intron Split Analysis results measuring relative changes in pre-mRNA (ΔIntron) and mRNA (ΔExon) levels from RNA-sequencing.

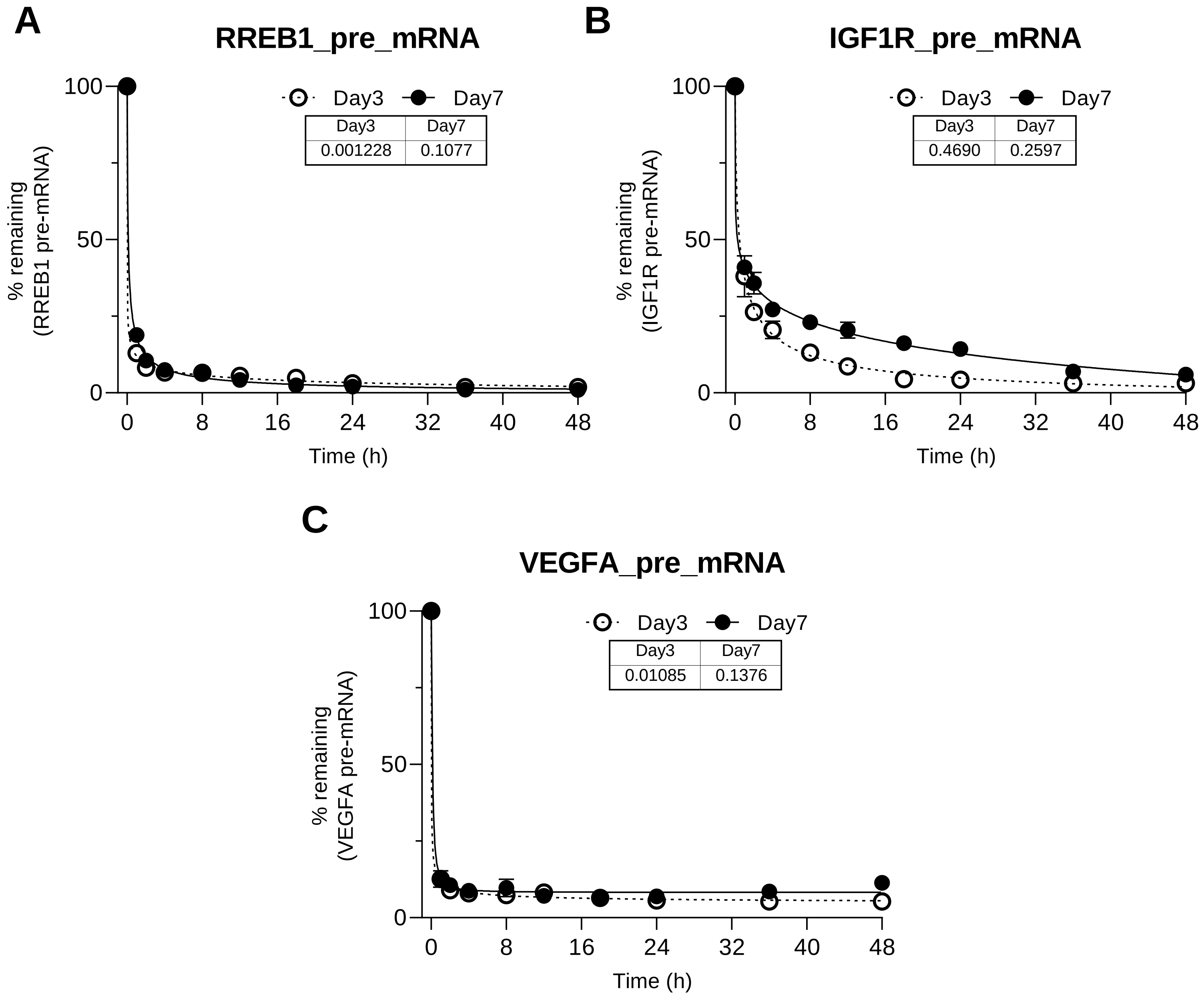


**Supplemental Figure 11.** (A-C) pre-mRNA stability assays for upregulated targets in RNA isolated from nuclear fraction from cells harvested on Day 3 (~50%) and Day 7 (~370%). RT-qPCR validating Actinomycin D treatment shut-down transcription at similar time after treatment in cells grown for 3 and 7 days. Transcription shut-down occurs within first hour independent of cell confluency (N=3).

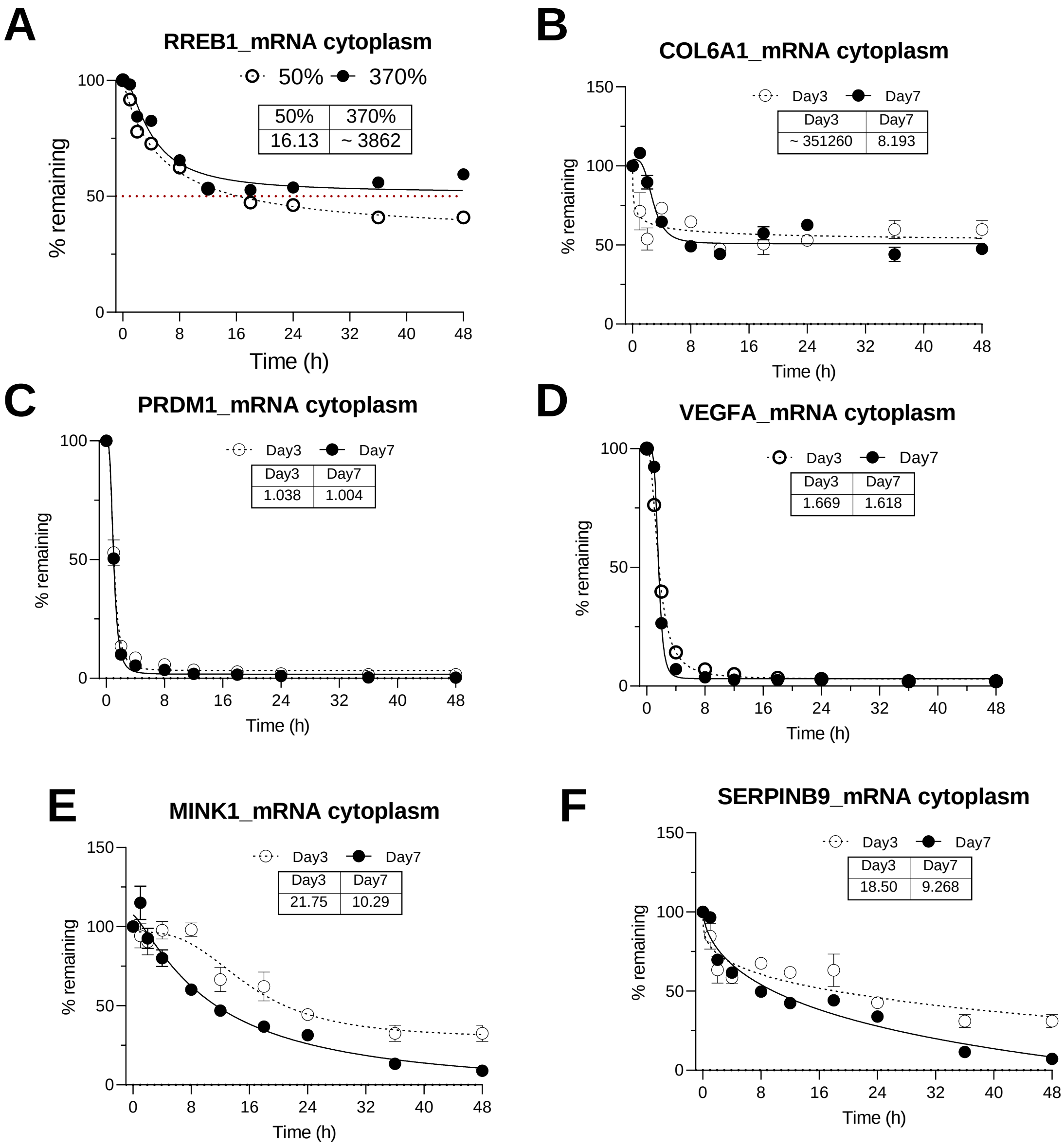


**Supplemental Figure 12.** (A-F) mRNA stability assays for upregulated targets in RNA isolated from cytoplasm fraction from cells harvested on Day 3 (50%) and Day 7 (370%). Table shows calculated estimate of half-life in hours on Day 3 and 7. (A-D) mRNAs had too long half-life or too short half-life to be compared or showed no difference between experimental conditions and (E-F) mRNAs showed longer half-life on Day 3 (50%).

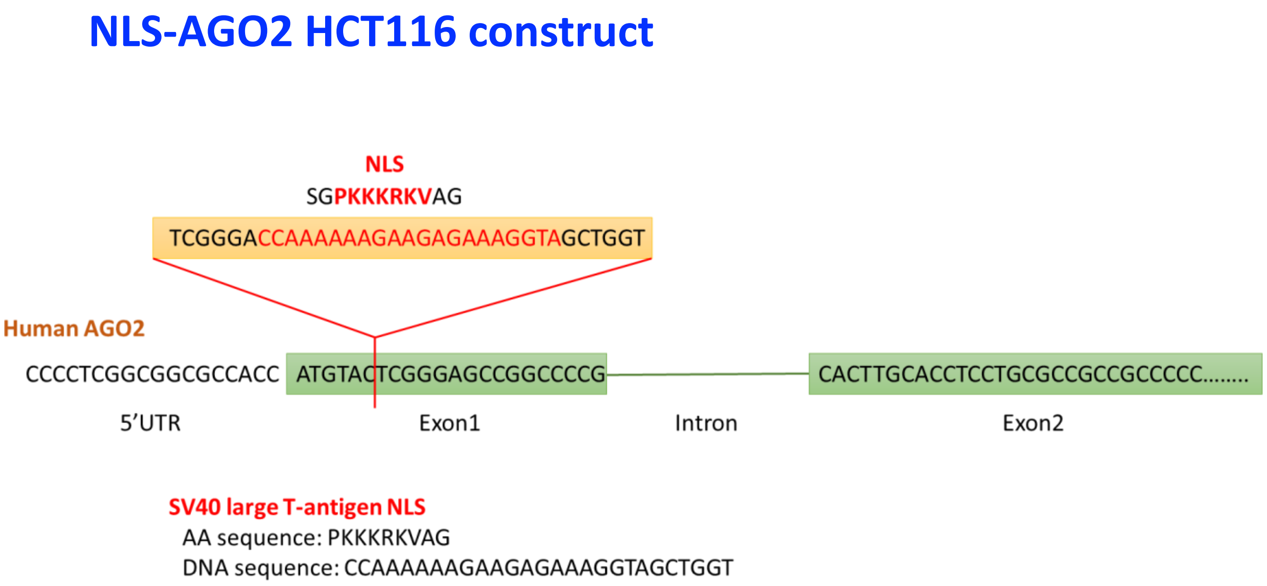


**Supplemental Figure 13. CRISPR/Cas9-mediated NLS-AGO2 HCT116.** SV40 NLS is inserted into N-terminal of human AGO2 according to previous papers. NLS fragment has 2 amino acid at both side as spacer sequences. NLS is inserted into both alleles to express AGO2 in only nucleus. To keep endogenous transcription level, endogenous promoter is used.

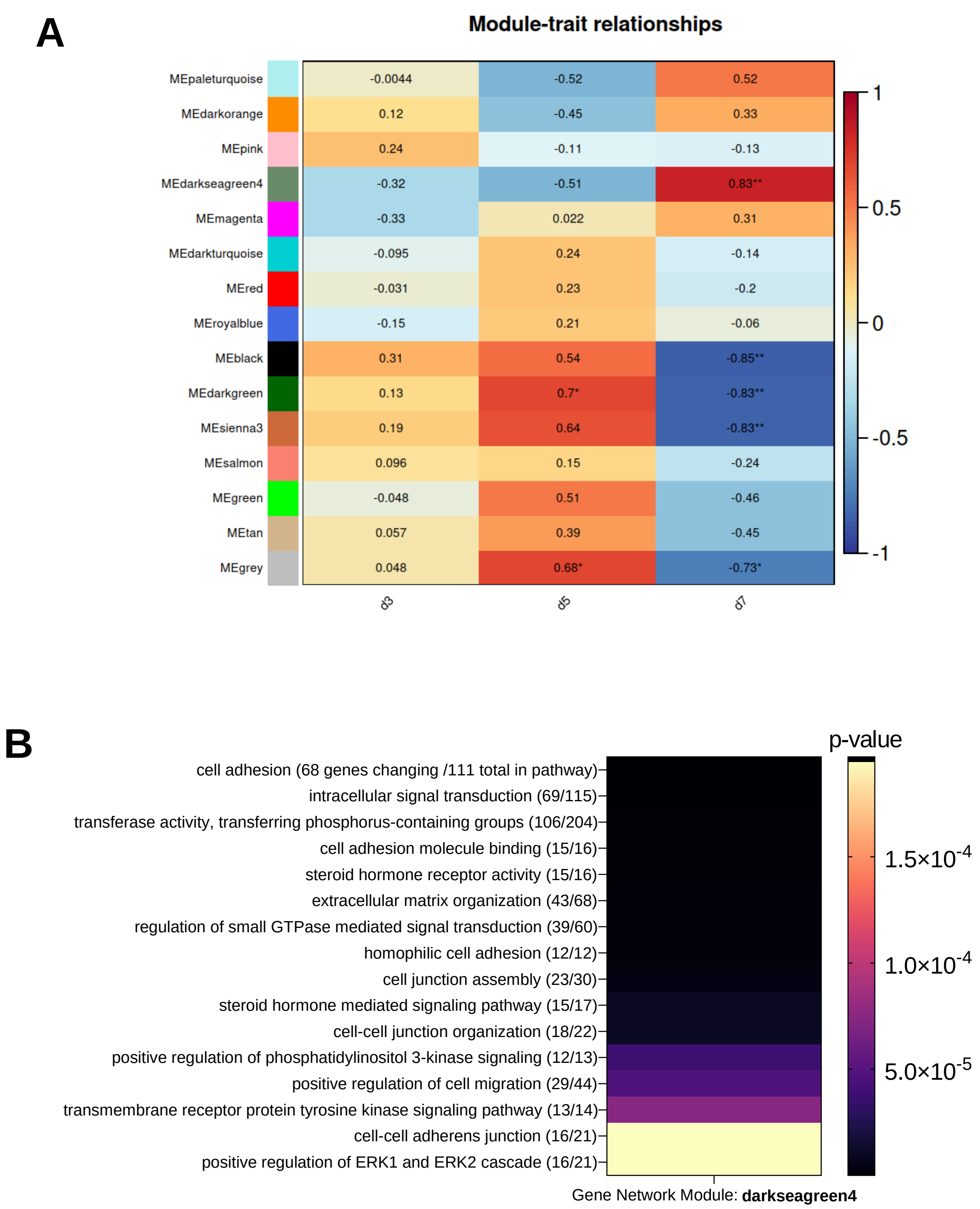


**Supplementary Figure 14.** (A) Results from Gene Network Analysis showing Pearson Correlation values for identifying groups/modules of genes named after different colors whose expression levels are positively or negatively correlated with different levels of cell density. D3 represents Day 3 at 50% confluency, D5 represents Day 5 at 250% confluency, and D7 represents Day 7 at 400% confluency. (B) Significance, p-value, of enriched pathways within "darkseagreen4" gene network module shown in (A). Ratio in parentheses show number of genes with differential expression change out of the total genes present in each pathway.

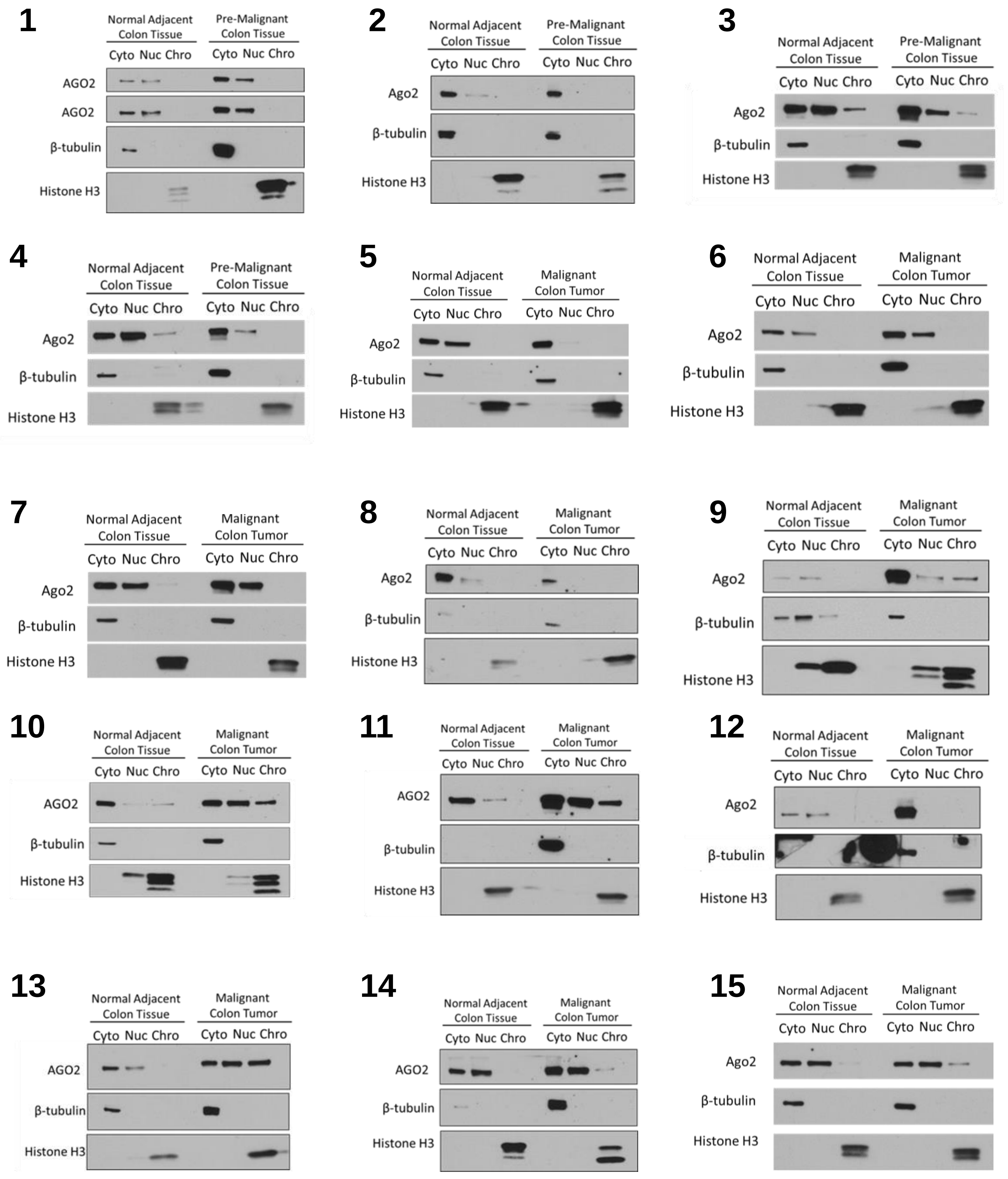
1
Normal Adjacent Colon Tissue
Pre-Malignant Colon Tissue
Cyto Nuc Chro Cyto Nuc Chro
AGO2
AGO2
β-tubulin
Histone H3
2
Normal Adjacent Colon Tissue
Pre-Malignant Colon Tissue
Cyto Nuc Chro Cyto Nuc Chro
Ago2
β-tubulin
Histone H3
3
Normal Adjacent Colon Tissue
Pre-Malignant Colon Tissue
Cyto Nuc Chro Cyto Nuc Chro
Ago2
β-tubulin
Histone H3
4
Normal Adjacent Colon Tissue
Pre-Malignant Colon Tissue
Cyto Nuc Chro Cyto Nuc Chro
Ago2
β-tubulin
Histone H3
5
Normal Adjacent Colon Tissue
Malignant Colon Tumor
Cyto Nuc Chro Cyto Nuc Chro
Ago2
β-tubulin
Histone H3
6
Normal Adjacent Colon Tissue
Malignant Colon Tumor
Cyto Nuc Chro Cyto Nuc Chro
Ago2
β-tubulin
Histone H3
7
Normal Adjacent Colon Tissue
Malignant Colon Tumor
Cyto Nuc Chro Cyto Nuc Chro
Ago2
β-tubulin
Histone H3
8
Normal Adjacent Colon Tissue
Malignant Colon Tumor
Cyto Nuc Chro Cyto Nuc Chro
Ago2
β-tubulin
Histone H3
9
Normal Adjacent Colon Tissue
Malignant Colon Tumor
Cyto Nuc Chro Cyto Nuc Chro
Ago2
β-tubulin
Histone H3
10
Normal Adjacent Colon Tissue
Malignant Colon Tumor
Cyto Nuc Chro Cyto Nuc Chro
AGO2
β-tubulin
Histone H3
11
Normal Adjacent Colon Tissue
Malignant Colon Tumor
Cyto Nuc Chro Cyto Nuc Chro
AGO2
β-tubulin
Histone H3
12
Normal Adjacent Colon Tissue
Malignant Colon Tumor
Cyto Nuc Chro Cyto Nuc Chro
Ago2
β-tubulin
Histone H3
13
Normal Adjacent Colon Tissue
Malignant Colon Tumor
Cyto Nuc Chro Cyto Nuc Chro
AGO2
β-tubulin
Histone H3
14
Normal Adjacent Colon Tissue
Malignant Colon Tumor
Cyto Nuc Chro Cyto Nuc Chro
AGO2
β-tubulin
Histone H3
15
Normal Adjacent Colon Tissue
Malignant Colon Tumor
Cyto Nuc Chro Cyto Nuc Chro
Ago2
β-tubulin
Histone H3

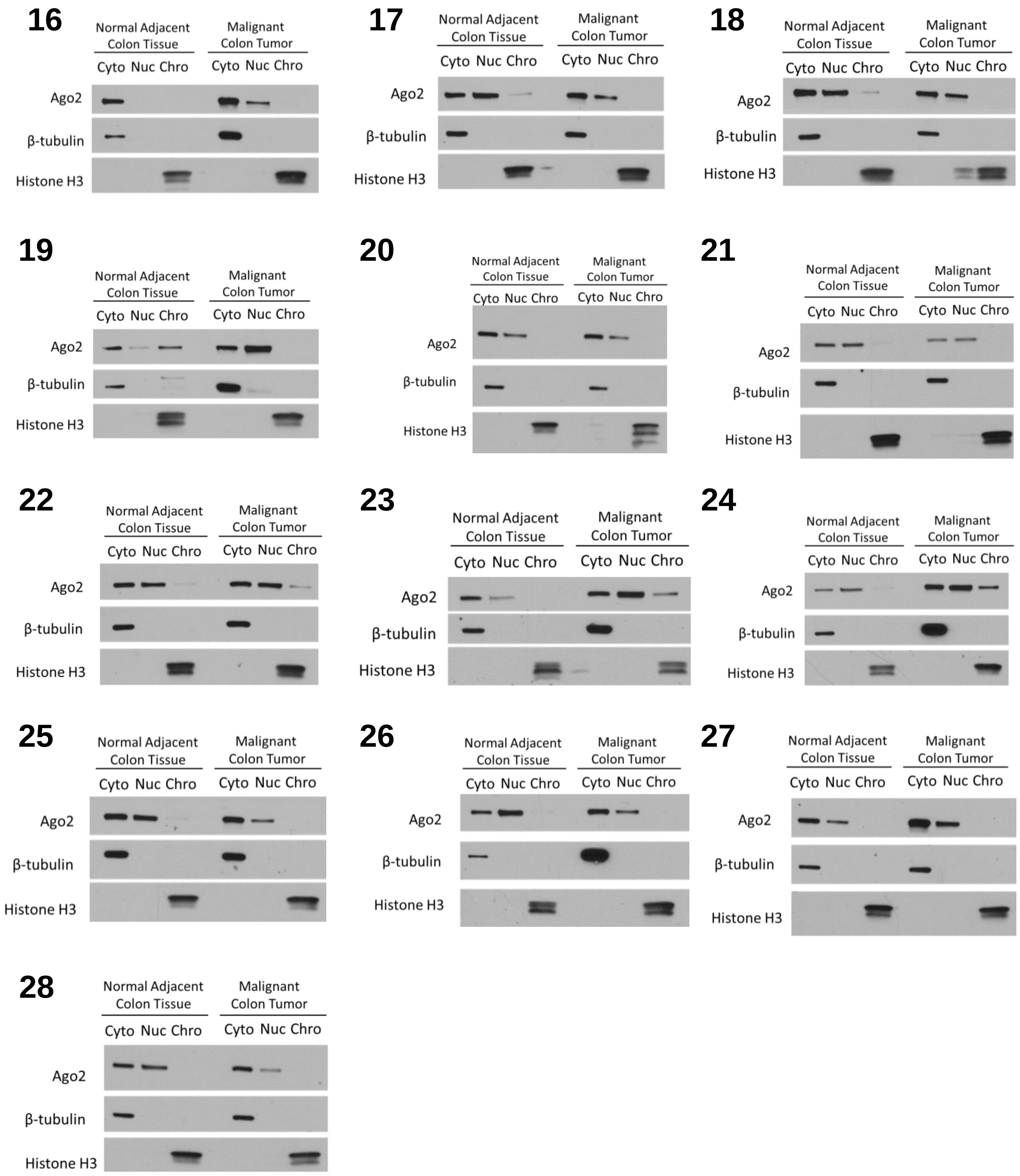


**Supplemental Figure 15. AGO2 Localization in Human Colon Tissue.**
(1-28) Western blot from pairs of tissue from Patients #1-28 fractionated into cytoplasm (cyto), soluble nuclear (nuc), and chromatin (chro) fractions. Pre-malignant indicates a colon adenoma tissue sample. Malignant indicates a colon adenocarcinoma tissue sample.

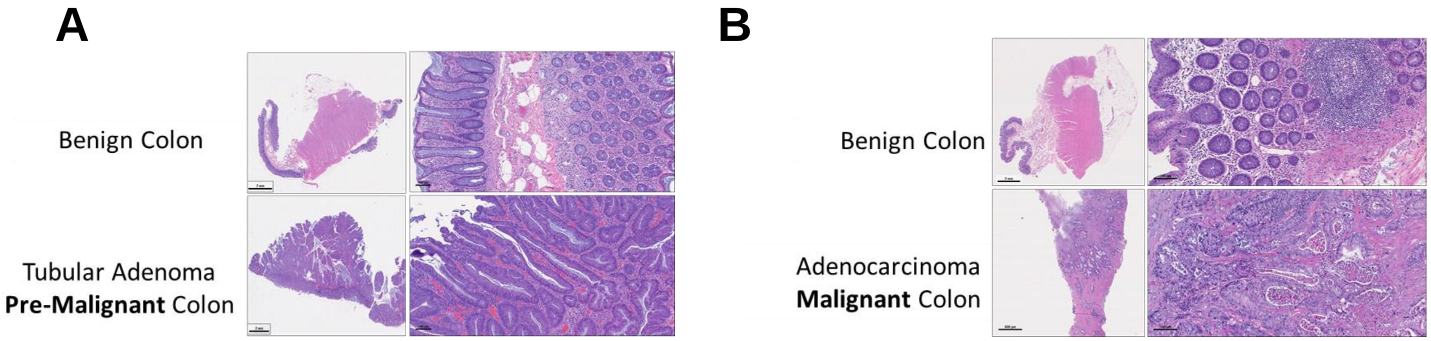


**Supplemental Figure 16.** (A, B) H&E staining of tissue slices from (A) Patient #2 and (B) Patient #10 used for pathology analysis.
